# Seasonal dynamics of a complex cheilostome bryozoan symbiosis – vertical transfer challenged

**DOI:** 10.1101/2022.12.09.519770

**Authors:** E.A. Bogdanov, A.E. Vishnyakov, O.N. Kotenko, A.V. Grischenko, A.V. Letarov, A.N. Ostrovsky

## Abstract

Symbiotic associations are dynamic systems influenced by both intrinsic and extrinsic factors. Here we describe for the first time the developmental and seasonal changes of the funicular bodies in the bryozoan *Dendrobeania fruticosa*, which are unique temporary organs of cheilostome bryozoans containing prokaryotic symbionts. Histological and ultrastructural studies showed that these organs undergo strong seasonal modification in the White Sea during the ice-free period. Initially (in June) they play a trophic function and support the development of a large population of bacteria. From June to September, both funicular bodies and bacteria show signs of degradation accompanied by development of presumed virus-like particles (VLPs); these self-organize to hollow spheres inside bacteria and are also detected outside of them. Although the destruction of bacteria coincides with the development of VLPs and spheres, the general picture differs considerably from the known instances of bacteriophagy in bryozoans. We broadly discuss potential routes of bacterial infection in Bryozoa and question the hypothesis of vertical transfer, which, although popular in the literature, is contradicted by molecular, morphological and ecological evidence.

## Introduction

Symbiotic associations are widespread among organisms ^1,2^, demonstrating an interorganismal level of morphological and physiological complexity in the framework of the holobiont paradigm ^3^. Given the huge variety of symbiotic associations — facultative and obligatory, commensal, parasitic and mutualistic — symbioses are also considered as the next level of adaptation of organisms to new types of biocenoses and competitive pressures ^4-6^, requiring an adjustment of the energy balance between the newly established symbiotic system and the environment ^7^.

The marine realm encompasses a plethora of symbiotic associations between various protists ^8,9^, metazoans ^10-12^, metazoans and protists ^13,14^, protists and prokaryotes ^15,16^, and metazoans and procaryotes ^17-19^. Some symbioses also involve viruses ^20^. While certain associations (e.g., corals-zooxanthellae, sponges-bacteria) became model study objects ^21-24^, most symbiotic relationships still await exploration. Among them are bryozoans hosting prokaryotes (reviewed in ^7,25^).

The phylum Bryozoa is composed almost exclusively of colonial animals inhabiting fresh, brackish, and salt waters from high to low latitudes and from the intertidal to the abyss. Bryozoans are suspension-feeders that mostly feed on unicellular algae, and, together with cnidarians and sponges, are among the dominating epibiotic groups in various benthic communities ^26-28^. A bryozoan colony consists of iterative modules termed zooids, usually less than 1 mm long. The feeding module (autozooid) is composed of a body wall (box-, tube- or sac-like cystid) and the retractile crown of ciliary tentacles associated with a U-shaped gut (polypide). Polypide excursions (extensions and retractions) are provided by the parietal and retractor muscles. Communication between zooids is provided via pores in zooidal walls, open or plugged by pore-cell complexes ^29-33^. In most marine bryozoans of the class Gymnolaemata, the zooids are additionally interconnected by a system of mesothelial funicular cords. These cross the zooidal cavity in different directions and transport substances throughout the colony ^34-37^.

Bryozoans form endosymbiotic associations with various prokaryotes that were first described by Lutaud ^38,39^. More recent studies have led to the conclusion that this association is a highly specialized symbiosis in which bacterial symbionts are vertically transmitted between generations ^40,41^, protect bryozoan larvae from predators ^42-45^, affect the fertility of colonies ^46,47^, and show signs of genome reduction ^48^. However, all of the experimental and molecular work behind this paradigm have focused on a small number of species from the genera *Bugula* (currently split into four genera) and *Watersipora*. Earlier microscopical studies showed the presence of prokaryotes (mostly intercellularly, but occasionally intracellularly) in species from nine phylogenetically distant families of cheilostome bryozoans (the only clade with published records of bryozoan symbionts). This indicates multiple independent origins of these associations ^38,39,49,50^ (reviewed in ^25^) and suggests various trajectories of bacterial circulation in the life cycle of bryozoan hosts.

In species of four bryozoan families (Bugulidae, Candidae, Beaniidae, Epistomiidae), the bacteria are inside the so-called funicular bodies (abbreviated as FBs here and elsewhere) – ‘capsules’ that have a wall of somatic cells associated with the funicular cords ^20,25,47,49,51^. It was suggested that these structures serve as sites for incubation and multiplication of bacteria ^25^, which can later move to the funicular cords and travel inside them towards incubation chambers containing a larva, thereby corroborating the idea of their vertical transfer ^47^. In addition, collapsing bacteria and a variety of virus-like particles were recently detected in the funicular bodies of two bryozoan species ^20^.

This paper presents ultrastructural details of co-specialization of the funicular bodies in the cheilostome bryozoan *Dendrobeania fruticosa* (Bugulidae) and its bacterial symbionts, along with putative virus-like particles (VLPs). We for the first time describe the ultrastructure of these temporary organs and their bacterial and viral content in their seasonal dynamics, considering the probability of bacteriophagy in this symbiotic system. We also discuss the structure, function, and development of the studied FBs, comparing them with published data. Importantly, we critically analyze the hypothesis of vertical transfer of symbionts in bryozoans and consider it and bacterial transmission from the external environment as possible alternatives for infection.

## Material and methods

The weakly-calcified erect colonies of *Dendrobeania fruticosa* ^52^ form delicate tufts 2–3 cm high with flat and narrow, dichotomously forking branches (Fig. 1A), anchored to the substrate by rhizoid-like polymorphs (kenozooids) ^37^. Each branch consists of 2–5, most frequently 3–4 rows of elongated autozooids. Some zooids are associated with the helmet-like brood chambers (ovicells) (Fig. 1B) bearing embryos during the reproductive season. In the White Sea, single embryos were recorded in June-August in the old parts of the colony, while active reproduction started in September-October, involving mostly the young colony parts.

**Figure 1.**
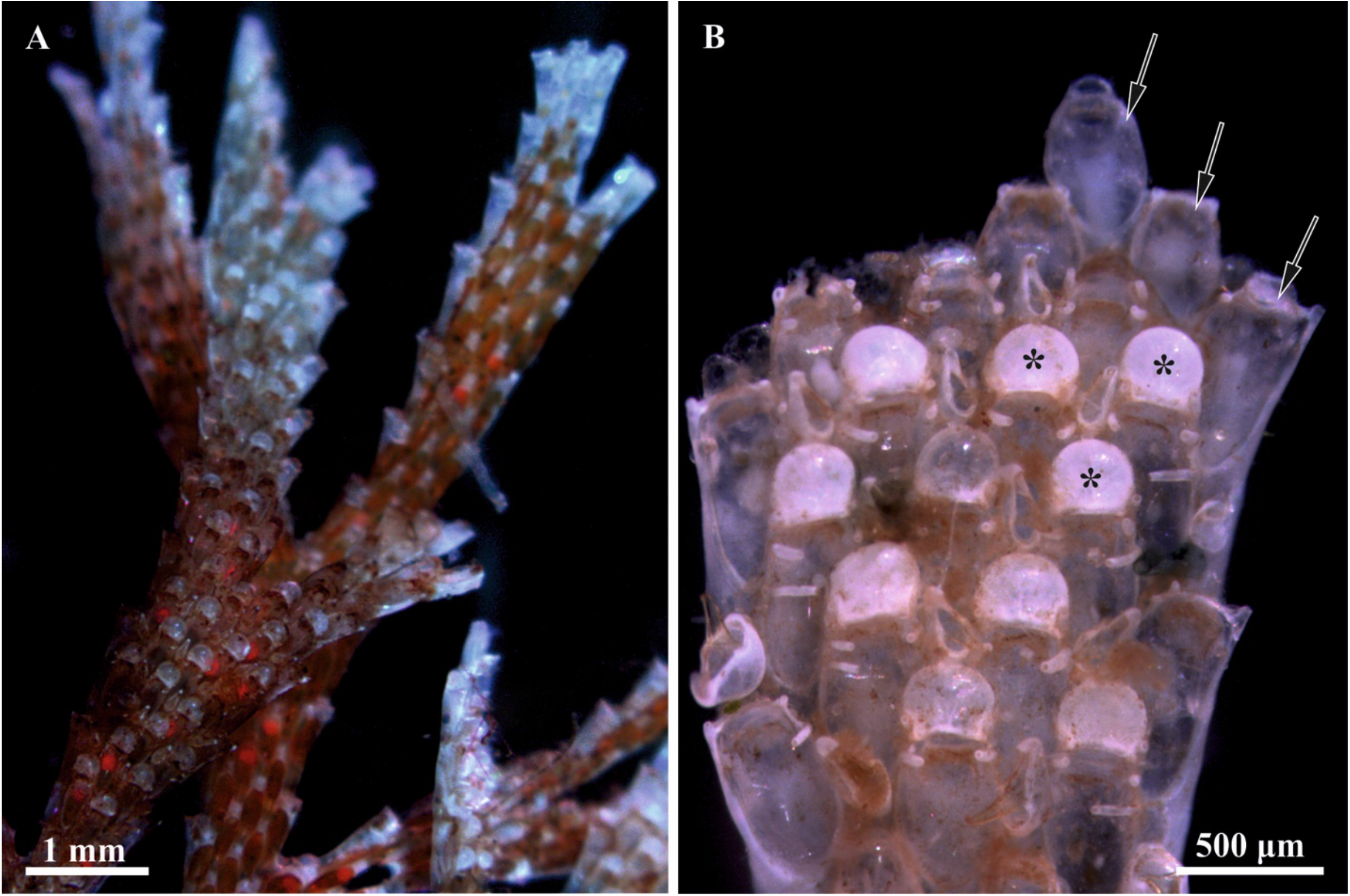
General view of a colony branch of *Dendrobeania fruticosa* with orange oocytes visible inside autozooids and embryos in ovicells (A) (collected on 23 August 2021), and colony fragment (B) (developing polypide buds inside young zooids on the tip of the branch shown by arrows, some ovicells marked by asterisks) (collected on 23 June 2021) (light microscopy).

*D. fruticosa* is a boreal-Arctic species, widely distributed in the seas of the northern hemisphere, from the Barents to Chukchi Sea and from the Beaufort Sea to Greenland; in the North Atlantic – from Iceland and the Norwegian Sea via the North Sea to Great Britain, and from the Saint Lawrence Gulf southwards to the Gulf of Maine; in the North Pacific – from the Bering Strait southwards to the northern part of the Sea of Japan, and from the Gulf of Alaska to Vancouver Island ^53-59^. This sublittoral to upper bathyal species has been recorded over a depth range 1.5–330 m, most frequently 7–25 m, on hard and mixed (with silt admixture) bottoms, at water temperatures of –1.9 to 14.5°C and salinities of 26.51–34.78‰. The substrata mainly include rocks, mollusc shells, hydroids, and other bryozoans.

Colonies of *D. fruticosa* growing on the lower part of boulders on the silty bottom were collected by SCUBA at 10—15 m depth on 21 March 2022, 14 June 2021, 22 June 2020, 23 June 2018, 19 August 2020, 23 August 2021 (one colony was taken in each case), and 31 September 2019 (three colonies), in the Chupa Inlet, Kandalaksha Bay, White Sea, close to the Educational and Research Station “Belomorskaia”, Saint Petersburg State University. No material was collected in July because of the weather conditions or logistical reasons. Colonies collected in September 2019 were kept at +4°C for two weeks before fixation. The others were fixed directly after collection.

For light microscopy and transmission electron microscopy (TEM), individual fragments of the colonies were fixed in 2.5% glutaraldehyde (buffered in 0.1 M Na-cacodylate with 10.26% sucrose, pH 7.4) for 1 hour, washed three times in buffer each lasting 15 minutes, and postfixed in 1% osmium tetroxide (OsO4) during 1 hour followed by three rinses in distilled water, each lasting 20 minutes. Samples were decalcified in 8.5% EDTA solution from a few hours to one day, after which the fragments were washed in distilled water. The dehydration process involved an ethanol series (30-50-70-80-90-100%) and acetone, after which the fragments were embedded in epoxy resin type TAAB 812. Semithin sections (1.0 μm thick) were made using a ultramicrotome Leica EM UC7 (Leica Microsystems, Wetzlar, Germany), stained by Richardson ^60^ and Humphrey & Pitman ^61^ techniques and examined with a light microscope Leica DM 2500. Ultrathin sections (70 nm thick) were also made using a Leica EM UC7 ultramicrotome. Sections were collected on copper grids and contrasted in uranyl acetate and lead citrate. The sections were examined using JEOL JEM-1400 and JEOL JEM-2100HC (JEOL Ltd., Japan) transmission electron microscopes and photographed with digital CCD cameras.

At least 26 zooids with FBs from nine colonies were studied ultrastructurally.

## Results

### Distribution and state of funicular bodies in the colony

All studied colonies of *Dendrobeania fruticosa* were overwintered. Funicular bodies (FB) were recorded in all of them (i.e. in all collection months: March, June, August and September). Most examined zooids contained a single FB in the cavity; in three zooids, two FB were found, and in one zooid, four bodies.

The state of the FBs (early and mature with no signs of degradation, and degrading at early, advanced and terminal stages) and their symbionts (‘healthy’ and modified) correlated with the month of collection and with the position and age of the zooids hosting them (for classification of the stages see Table 1 and Fig. 2). Early and mature FBs with no signs of degradation were recorded in young zooids in the colonies collected only in June, whereas older zooids in the colonies collected in June and August contained FBs (and bacteria) in various stages of degradation, from the ‘early-’ to ‘late-advanced’. Colonies collected in late March and late September contained only strongly degraded FBs in old zooids.

**Table 1.** State of the funicular bodies (FB) and bacteria, and the presence of the virus-like particles (VLP) in the colonies of *Dendrobeania fruticosa* collected in different months.

| State of funicular body | early and mature functional FBs with no signs of degradation | initial stage of degradation | -advanced stage of degradation |  |  | terminal stage |
| --- | --- | --- | --- | --- | --- | --- |
|  |  |  | early- | mid- | late- |  |
| % of bacteria | 100% | 100% | 100-70% | 70%-30% | 30-1% | 1-0% |
| State of bacteria | ‘healthy’ | ‘healthy’ or slightly modified | modified. VLP may be present | modified VLP often present | modified, VLP mostly present<br>spheres and fibrils often present | modified |
| Inner cells | non-modified microvilli present. | slightly modified microvilli mostly lost. | modified | modified often flattened | modified mostly flattened | mostly double membrane present |
| Interlayer space | absent | absent or small, with small amount of VLP inside | small | large, densely filled with VLP | spherical complexes may be present | spherical complexes may be present |
| Collection date | 23 June 2018<br>22 June 2020 | 23 June 2018 | 14 June 2021 | 19 August 2020<br>23 August 2021 | 14 June 2021<br>23 August 2021<br>31 September 2019 | 14 June 2021<br>21 March 2022 |

**Figure 2.**
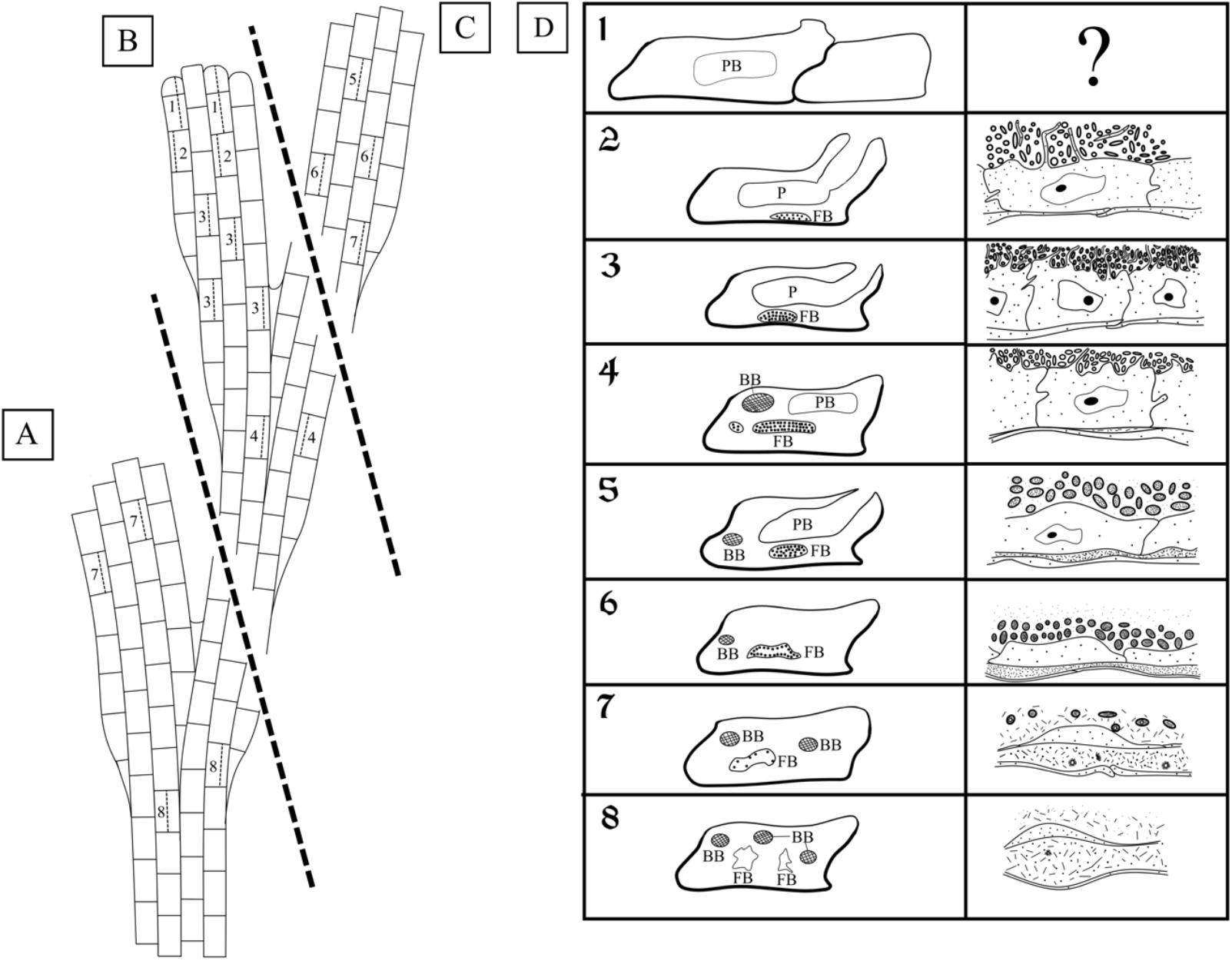
Schemes of fragments of colonies collected in June (B), June-August (C) and September (A), and showing position of autozooids containing FBs at different stages of development and degradation. In A and C distal tips of the branches are not growing. Distal tip of the fragment in B shows zooidal buds, indicating a growing branch. D, Vertical columns show schematic depictions of longitudinal sections of autozooids (left) and partial FBs (right) at consecutive stages of development (1-8) and degradation (corresponding to Table 1): 1– distal zooidal bud and young autozooid with a polypide bud inside (no FBs); 2 – young autozooid with functional polypide and early developing FB; 3 – young autozooid with functional polypide and mature FB; 4 –autozooid with brown body, polypide bud and mature FB at initial stage of degradation; 5 – autozooid with brown body, functional polypide and FB at early-advanced stage of degradation; 6 – autozooid with brown body and FB at mid-advanced stage of degradation; 7 – autozooid with brown bodies and FB at late-advanced stage of degradation; 8 – autozooid with brown bodies and two FBs at terminal stage of degradation. Abbreviations: bb, brown body; fb, funicular body; p, polypide; pb, polypide bud.

In the colony collected on 23 June 2018, the growing branch tip included a terminal zooidal bud, proximally followed by a young zooid with a polypide bud (Fig. 1B), and then two young zooids with functional polypides. In none of these four modules were FBs observed. In contrast, the following (towards the base of the branch) six zooids in this row had functional polypides and FBs from the early developing (in two distalmost zooids among those six) to the non-modified mature FBs (in the following four zooids). None of the FBs showed any signs of degradation and contained numerous and ‘healthy’ (non-modified) bacteria (Figs 3C, 4–6). We also examined sections of a zooid (presumably the most proximal module in this row) with degenerating polypide and mature non-modified FB (Fig. 3A). Thus, this colony piece showed an entire sequence from the early to mature stages in the FBs and zooidal development. Also, in the tip of another branch the youngest FB was recorded in the third zooid from top: the first was a zooidal bud, the second was a young zooid with a polypide bud, and the third was a zooid with fully-formed polypide and small FB.

**Figure 3.**
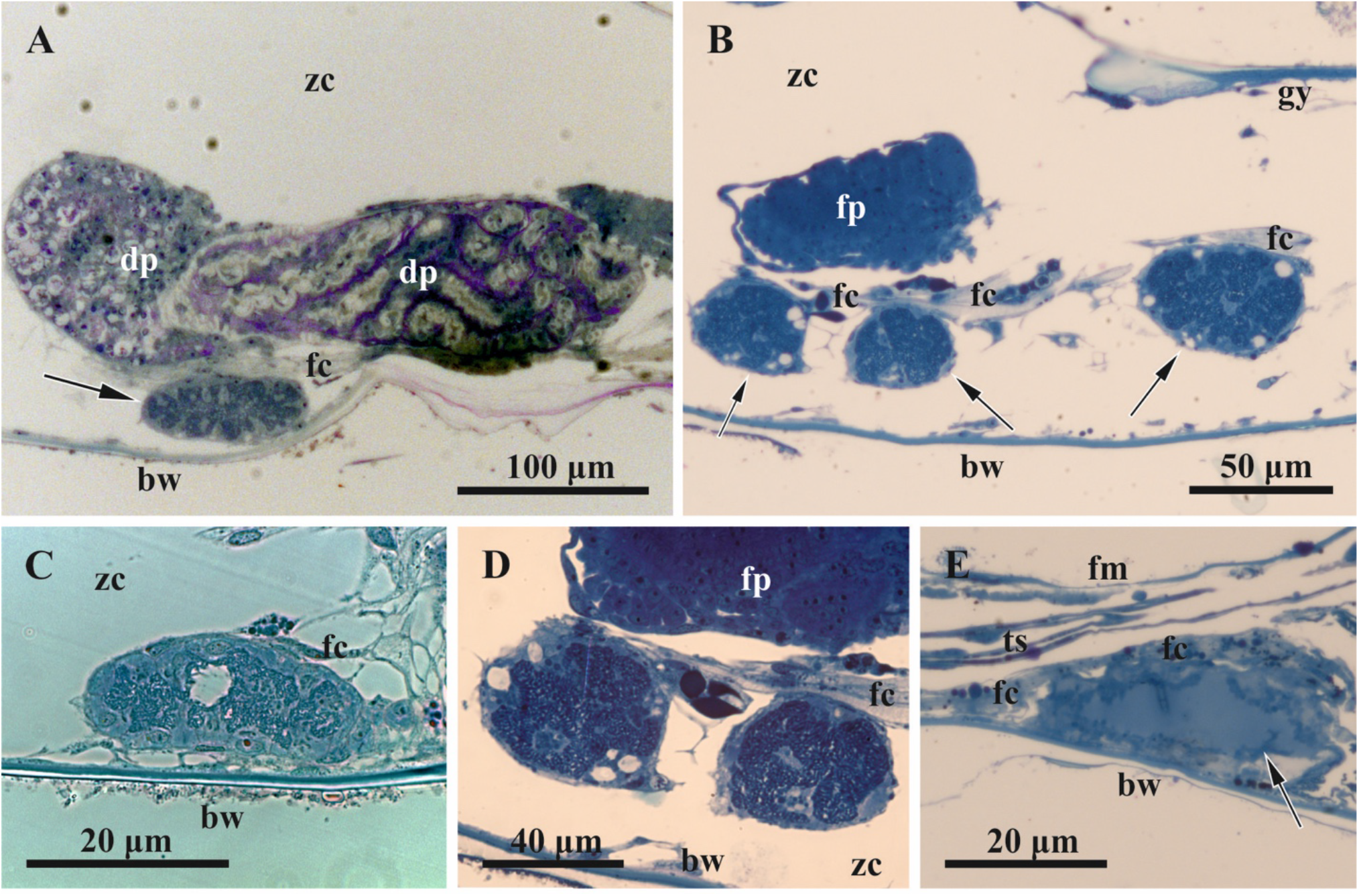
General view of funicular bodies inside the zooidal cavity in *Dendrobeania fruticosa* (A-D, collected on 23 June 2018, E, collected on 31 September 2019) (longitudinal stained sections, light microscopy). A, Mature non-modified FB (arrow) situated between the basal zooidal wall and the degenerating polypide; part of the funicular cord is sandwiched between the FB and the polypide. B, Zooid showing two cross-sectioned parts of one lobed FB below growing polypide bud, and part of the second FB to the right (all FBs shown by arrows); a large funicular cord runs above both FBs. C, Mature non-modified FB situated on basal wall of zooid and connected to thin processes of funicular cells. D, Enlarged view of two parts of one FB shown in B (both FBs show the initial stage of degradation, visible as large ‘vacuoles’ potentially reflecting either cell degradation or a fixation artifact). E, Collapsing FT. Abbreviations: bw, basal wall; dp, degenerating polypide; fc, funicular cord; fm, frontal membrane; fp, forming polypide bud; gy, gymnocyst; ts, tentacle sheath; zc, zooidal cavity.

**Figure 4.**
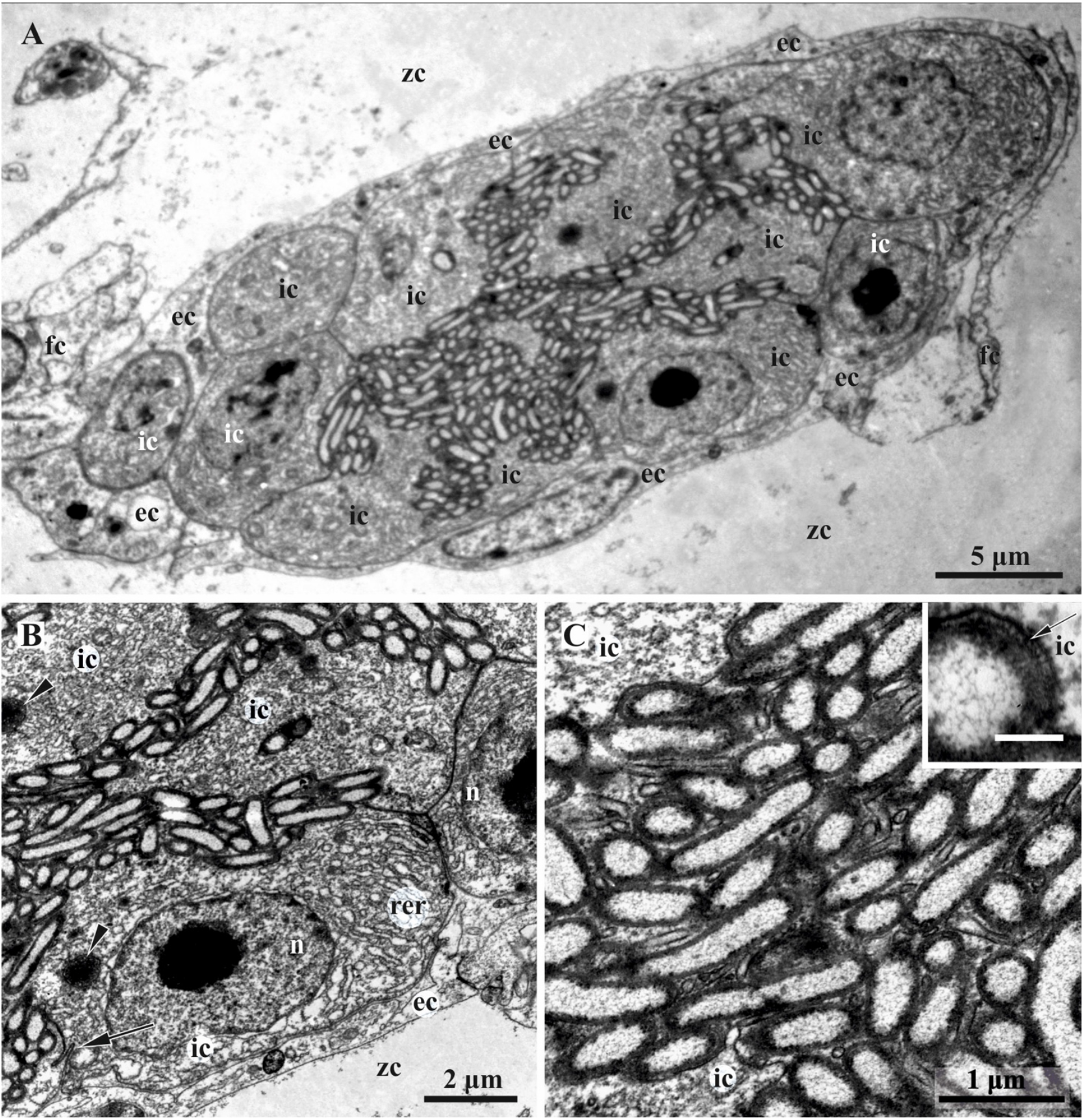
Early funicular body of *Dendrobeania fruticosa* (collected on 23 June 2018) (TEM). A, Whole view of early-stage funicular body. B, Partial view of cavity and wall of early-stage FB showing external and inner cell layers (arrowheads: multivesicular bodies; arrow: adherens junction with Z-curve between cells). C, Bacterial symbionts and microvilli formed by cells of inner layer. Inset: enlarged area showing bacterial pili (arrow). Abbreviations: ec, cells of external layer; fc, cells of funicular cords; ic, cells of inner layer; n, nucleus; rer, rough endoplasmic reticulum; zc, zooidal cavity.

Older zooids in the basal part of the same branch contained FBs showing the first signs of degradation (initial stage) and bearing numerous non- or slightly modified bacteria (Figs 7, 10A). These zooids also contained a brown body (remnants of a degenerated polypide), and one of them had a new polypide bud (Fig. 3B, D), whereas another seemed to hold a degenerating polypide.

In a growing tip of the colony collected on 22 June 2020, a young zooid with functional polypide contained the largest recorded mature FB that was non-modified, and enveloped numerous ‘healthy’ bacteria.

The colony collected on 14 June 2021 contained FBs at different stages of degradation. In the distal colony part, young zooids with functional polypides forming apical and lateral new growing tips of the branches contained FBs in the ‘early-advanced’ stage of degradation. These FBs still enveloped numerous bacteria. Old zooids without polypides and with brown bodies forming the apical non-growing tip of the studied branch contained ‘late-advanced’ FBs, either shrunken or swollen, with few bacteria. In the basal part of the same colony, old zooids (without polypides and with fragments of brown bodies) forming a branch near the zone of rhizoids, contained large swollen FBs at the terminal stage of degradation and almost no bacteria. FBs in the two latter groups of old zooids are probably overwintered (i.e. developed in the previous year). Similarly, in the colony collected on 21 March 2022, old zooids without polypides in the apical, obviously overwintered non-growing parts of the branches had FBs at the terminal stage of degradation.

In the colonies collected in the second half of August, zooids forming the apical non-growing parts of the branches contained FBs at either ‘mid-advanced’ (19 August 2020 and 23 August 2021) or ‘late-advanced’ (23 August 2021) stages of degradation. Bacteria were still common in some of them (‘mid-advanced’), although most FBs had only a few (Figs 8A, 10B). Zooids with FBs may or may not have a functioning polypide, also possessing a brown body.

In the colonies collected on 31 September 2019, a few symbionts were detected in the FBs. They were at the ‘late-advanced’ stage of degradation (Figs 3E, 8B, C, 10C). Examined zooids in the apical non-growing parts of the branches mostly lacked polypides.

Typically, FBs were adjacent to the basal wall of the zooid (Figs 3A, C, E, 6A), although some were suspended inside the zooidal cavity underneath the forming (Fig. 2B), degrading (Fig. 3A) or functional (Fig. 3D) polypide. Regardless of the position, FBs were ‘anchored’ by either large funicular cord(s) or by thin processes of their cells, or often by both (Figs 3—5, 8B, 10). Some funicular cords showed small internal lacunae, but they were never seen to be connected with the internal cavity of FB in our TEMs.

**Figure 5.**
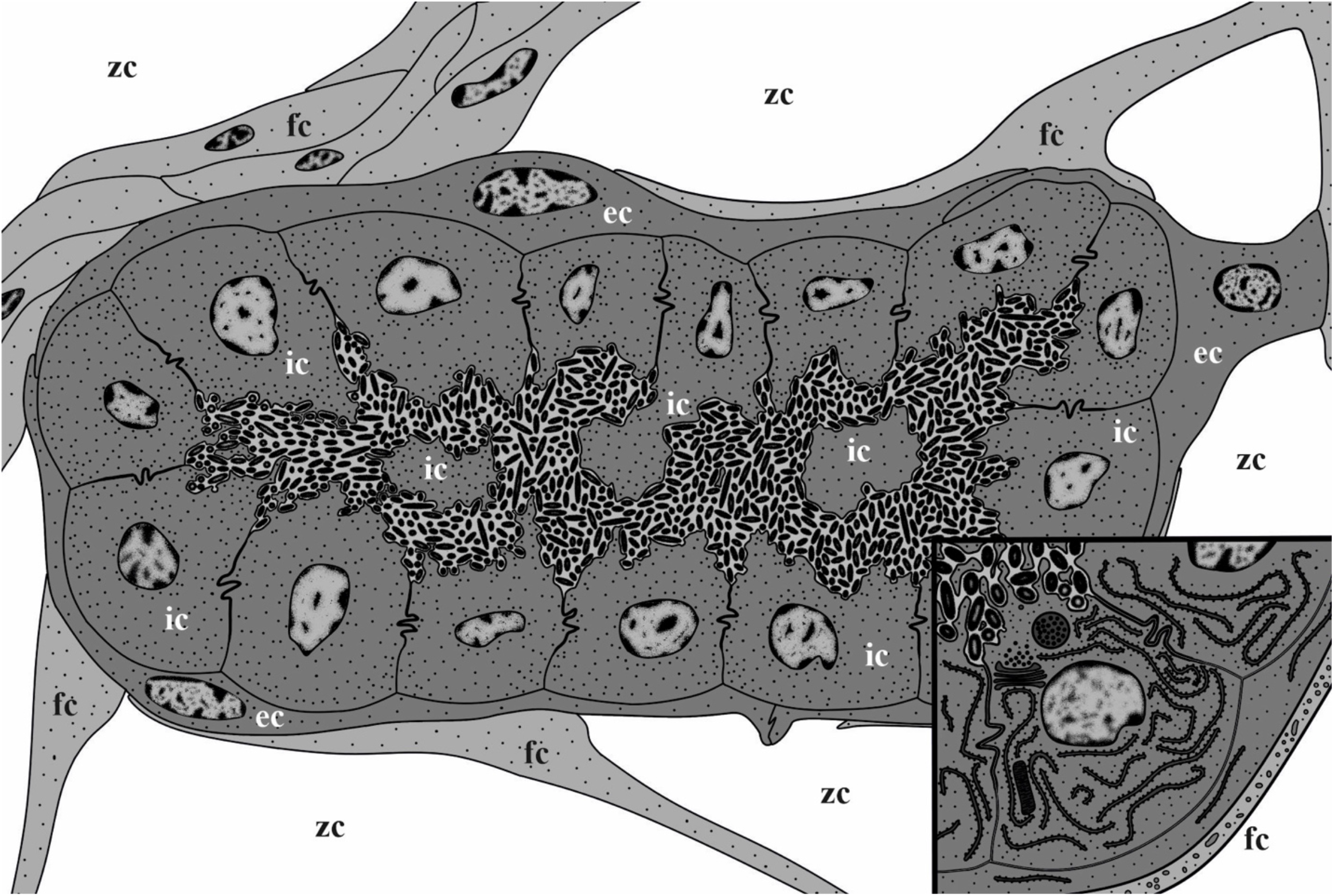
Scheme of a mature non-modified funicular body of *Dendrobeania fruticosa* (corresponding to those collected on 23 June 2018) showing two-layered wall structure, thick funicular cord (to the left), and thin processes of funicular cells adjacent to the external cell layer, and bacteria inside the FB internal cavity. Inset: enlarged area showing cells of both external and inner layers and bacteria with cytoplasmic processes in between (not shown in the larger scheme). Abbreviations: ec, cells of external layer; fc, cells of funicular cord; ic, cells of inner layer; zc, zooidal cavity.

### Early and mature non-modified funicular bodies (June)

All but one young zooid with early and mature non-modified FBs collected in June 2018 and 2020 had the first functional polypide; the exception was one zooid in which the polypide had begun to degenerate (Fig. 3A). The length of these FBs varied from 40 to 180 μm, the diameters from 15 to 50 μm.

Early and mature FBs had an oval or elongate-oval shape and consisted of a wall of somatic cells and an internal cavity filled with bacteria (Fig. 3A, C). In early FBs, this cavity was slit-like and ‘branching’ (Fig. 4A), in mature FBs it was much larger (Figs 3A, C, 5, 6A) and divided into many connected pockets.

**Figure 6.**
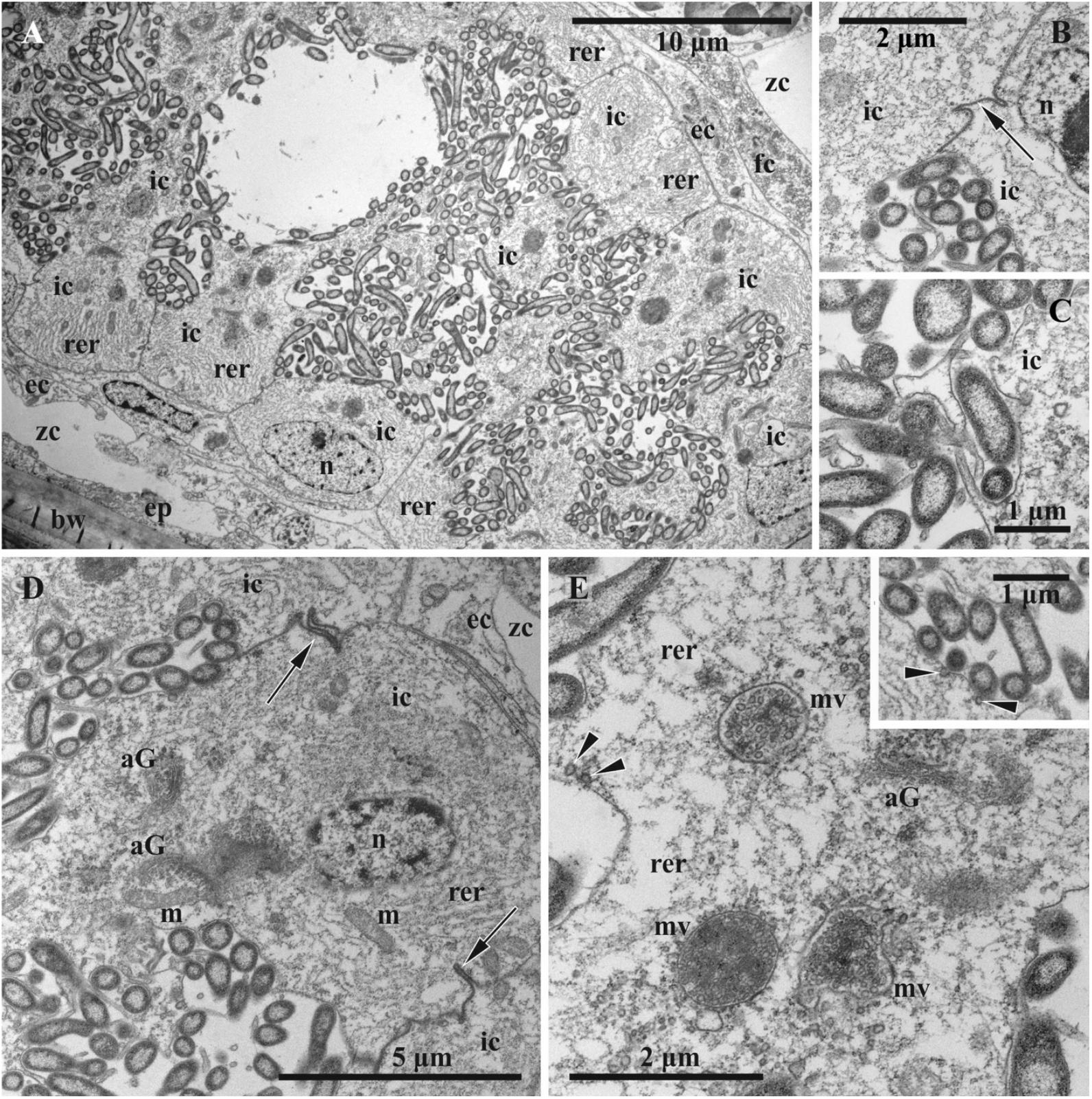
Ultrastructure of mature non-modified funicular body of *Dendrobeania fruticosa* (collected on 23 June 2018, also shown in Fig. 3C) (TEM). A, Part of FB with two-layered wall and adjacent funicular cells; non-modified bacteria fill the internal space. B, C, Area of internal cavity with bacterial symbionts and microvilli formed by cells of inner layer. D, E, Details of ultrastructure of cells of inner layer; extensive RER, Golgi apparatuses and large multivesicular bodies in various stages of development are visible (inset and E: presumed exocytosis shown with arrowheads) (arrow: adherens junction with Z-curve between cells of inner layer in B and D). Abbreviations: aG, Golgi apparatus; ec, cells of external layer; ep, epithelium of the body wall; fc, cells of funicular cords; ic, cells of inner layer; m, mitochondrion; mv, multivesicular body; n, nucleus; rer, rough endoplasmic reticulum; zc, zooidal cavity.

A TEM study showed that the wall of both early and mature FBs is formed by two cell layers of contrasting ultrastructure (Figs 4—6). The external (outer) flattened cells were arranged in a single layer and sometimes overlap each other. They had an electron-translucent cytoplasm with a few organelles (elongated nucleus with prominent peripheral heterochromatin, sparse cisternae of rough cytoplasmic reticulum (RER), and some microvesicles). They resembled the funicular cells directly contacting them and sometimes gave the impression of the presence of a third cell layer (Figs 4A, 5, inset, 6A). Overall, the external cells showed little evidence of synthetic or transport activity.

The inner layer consisted of cuboidal and prismatic cells, some having long irregular outgrowths separating a FB cavity onto interconnected chambers/pockets densely filled with bacteria (Figs 3A, B, 5, 6A). These cells had a distinct polarity: their basal membrane abutted the membrane of the cells of the outer layer, whereas the apical one formed multiple thin cytoplasmic processes (microvilli) protruding between bacteria (Figs 4B, C, 5 and inset, 6A—D). The lateral surfaces of the inner cells have tight contacts exhibiting a Z- or V-shaped configuration (Figs 5, 6B, D). The cytoplasm was electron-translucent, although denser than in the external cells, and contained a large nucleus, mostly filled with euchromatin and highly fragmented heterochromatin, an extensive RER, several Golgi complexes, and abundant multivesicular bodies (Figs 4—6). Microvesicles and pits were detected beneath and in association with the apical membrane, possibly indicating exo- and endocytosis (Fig. 6E and inset).

The symbionts filling the FB cavity were Gram-negative, elongate-oval or rod-shaped bacteria 2-3 μm long, some up to 5 μm, and about 0.5 μm in diameter (Figs 4, 6). A well-defined, central, electron-lucent nucleoid zone with flocculent contents was surrounded by a thin peripheral layer of electron-dense cytoplasm, enveloped by two membranes. Bacterial cells (some obviously dividing) were in contact with the apical membrane of the inner cells of FT and sometimes clearly showed pili on their surface (Fig. 4C, inset). The space of the FB cavity between bacteria was electron-transparent. No connection between this cavity and the lacunae of funicular cords was detected.

### Degrading funicular bodies (June, August, September)

#### Initial stage

In the colony collected on 23 June 2018, zooids in the basal part of the growing branch contained FBs in the initial stage of degradation (Table 1, see also above). Usually they were elongate-oval, in two cases lobed. Three studied autozooids contained two FBs each (in once case an oval and a bilobed FB occurred in the same zooid), and one zooid contained four FBs (one of them with four lobes, partly shown in Fig. 3B, D, see also Fig. 7). The latter zooid had a new polypide bud, and all these zooids contained a brown body.

**Figure 7.**
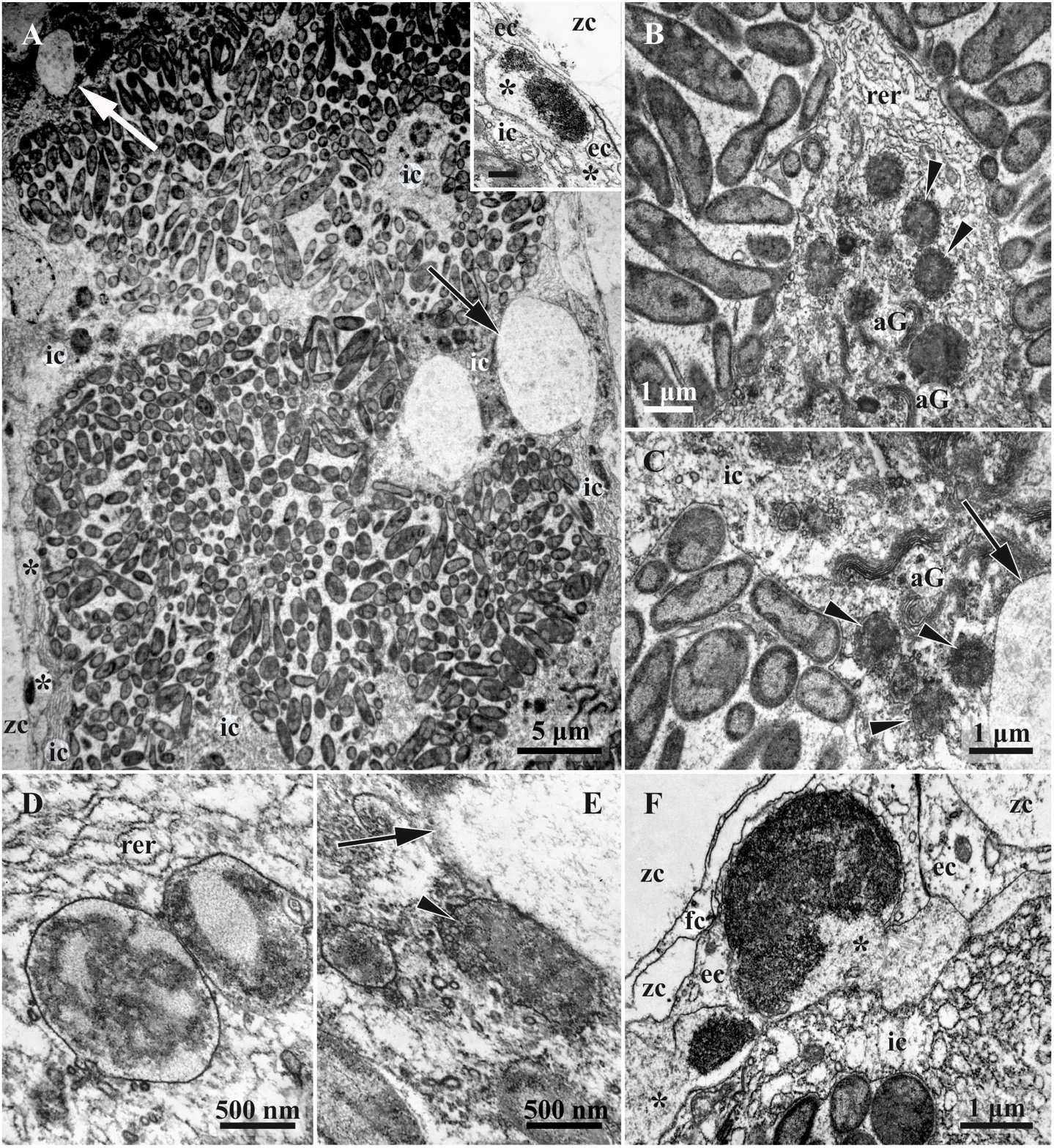
Ultrastructure of funicular bodies of *Dendrobeania fruticosa* (collected on 23 June 2018) showing the initial stage of degradation (TEM). A, Part of FB densely filled with modified bacteria; large expanded parts of RER shown by arrows (also visible in C and E) (inset: peripheral area of FB wall showing cells of both external and inner layers and forming ‘interlayer’ space (asterisk) containing electron-dense bodies; scale bar 500 nm). B, C, Bacteria and cells of inner layer with presumed phagosomes (arrowheads) containing bacteria-like content. D, E, Presumed phagosomes (shown by arrowhead in E) with bacteria-like content shown at higher magnification. F, Peripheral area of FB wall showing cells of external and inner layers with an interlayer space (ILS) (asterisk) between them containing electron-dense bodies and ‘fibrils’ sometimes in groups (arrowhead) (also visible in inset). Abbreviations: aG, Golgi apparatus; ec, cells of external layer; fc, funicular cell; ic, cells of inner layer; rer, rough endoplasmic reticulum; zc, zooidal cavity.

At this earliest stage of FB modification, cells of the external layer were unchanged. In the cells of the inner layer, RER was still massive, but some cisternae formed local expansions, sometimes turning into large ‘vacuoles’ and occupying up to a third of the cell volume (Fig. 7A, C, E, and possibly 3B, D). Cisternae of the Golgi complexes became longer and formed dense piles. In addition to multivesicular bodies, phagosomes with heterogeneous contents, which in some cases resemble lysing bacteria, appeared in the cytoplasm (Fig. 7B-E). Initially consisting of several non-connected lacunae, an interlayer space (ILS) was formed between cells of the external and inner layers of FB. It was mostly narrow, becoming wider in places containing large electron-dense bodies of various sizes (Fig. 7A, inset, F). In addition, tiny fibrils about 100 nm long and angular globules (presumably virus-like particles, see below) became visible in some areas of ILS (Fig. 7F).

At the initial degradation stage, numerous bacteria, some apparently still dividing, filled the entire cavity of the FBs (Figs 7A, 8A). However, the number of microvilli formed by the inner cells and present between bacteria, was significantly reduced. Microvesicles beneath the apical membrane of these cells were abundant, presumably indicating transmembrane transport. The morphology of bacteria changed too. Although the maximum length was still about 5 μm, many bacteria were about 1 μm in diameter. The most noticeable changes occurred in the bacterial nucleoid, which was fragmented and became smaller (Fig. 7A-C).

**Figure 8.**
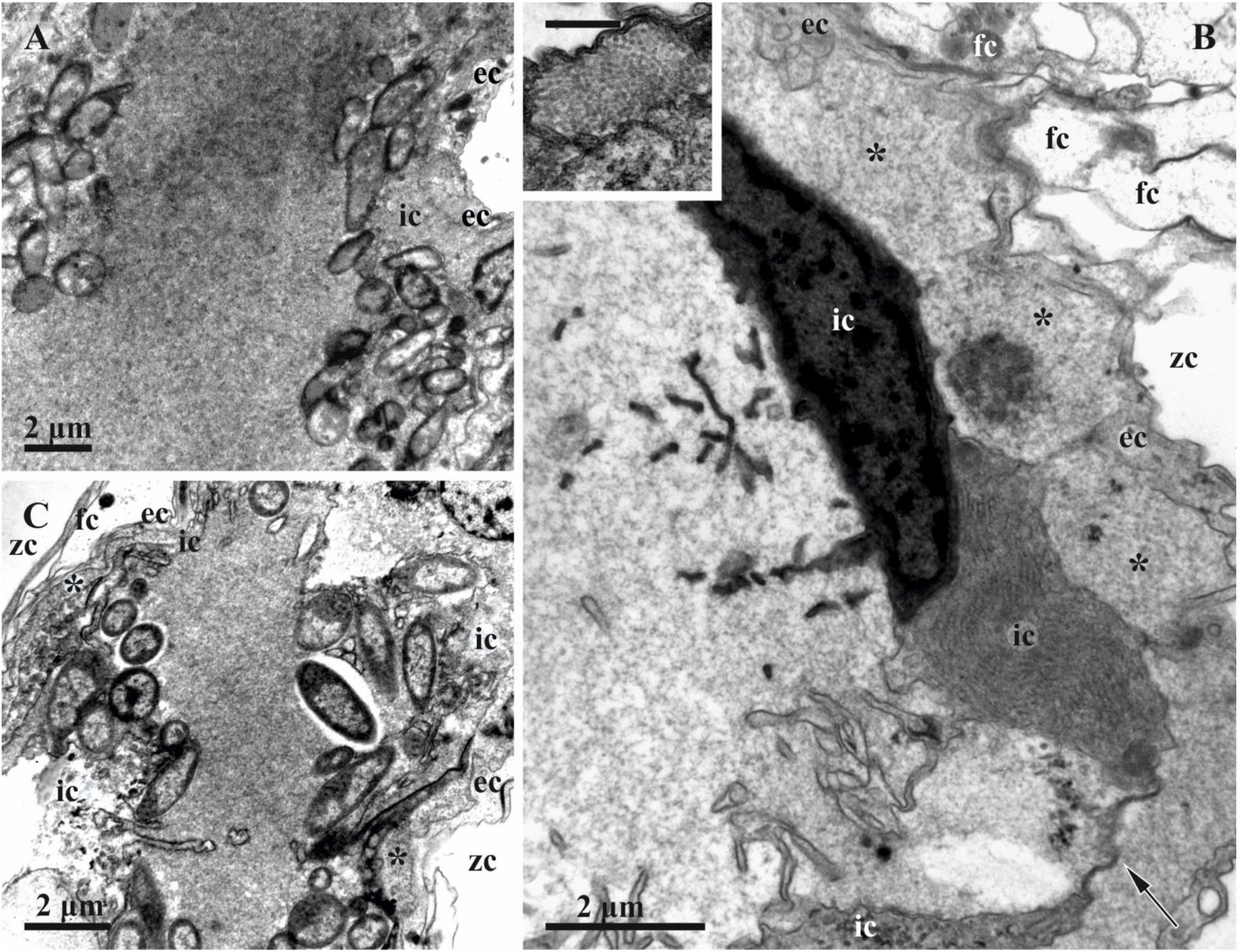
Degradation stages of the funicular bodies and their bacterial symbionts in *Dendrobeania fruticosa* (TEM). A, Middle-advanced stage showing remaining bacteria inside the cavity of FB near its inner cells; peripheral part of FB with two cell layers and ILS (asterisk here and elsewhere) inbetween are visible in the right upper corner (19 August 2020). B, Area of FB at the late-advanced stage showing cavity devoid of bacteria and its wall consisting of inner and external cell layers with greatly expanded ILS; inner layer consists of collapsing cells with cytoplasm of various electron density, some still having long and branching cytoplasmic processes; ‘double’ membrane (arrow) – part of collapsing cell, is visible between two cells (inset: enlarged fragment of ILS with tightly packed ‘globules’; scale bar 500 nm) (31 September 2019). C, Area of FB at the late-advanced stage of FB degradation showing remaining bacteria collapsing inside FB cavity (31 September 2019). Abbreviations: ec, cells of external layer; fc, funicular cell; ic, cells of inner layer; zc, zooidal cavity.

#### Advanced stages

In the colony collected on 14 June 2021, FBs were either at the early, or late-advanced, or even at the terminal stages of degradation, depending on zooid age (Table 1, see above). Middle-advanced stages were detected in the colonies collected on 19 August 2020 and 23 August 2021. Late-advanced stages were recorded in the colonies collected on 23 August 2021 and 31 September 2019. In zooids with advanced-stage FBs, functional polypides were usually absent, being substituted by their remnants in the form of brown bodies; nonetheless, even at the end of September, we found several zooids with such FBs and with functional polypides (Fig. 3E).

At the early-advanced stage of degradation, FB structure was similar to the initial stage, although the number of bacteria was markedly reduced and their morphology changed. The interlayer space of FBs was more developed and sometimes contained virus-like particles (Fig. 10A).

The middle, late and terminal stages of FB degradation differed drastically from the initial and early-advanced stages. E.g. in longitudinal semi-thin sections, some FBs at the late-advanced stage were unevenly elongated, either shrunken or swollen, with a dark periphery and lighter homogeneous contents (Fig. 3E). The bacterial cells inside them were almost indistinguishable with light microscopy. The length of such FBs could reach 100 μm, and the width varied within 10-30 μm due to numerous deformations (Figs 3E, 10B, C).

On TEM images, the outer cell layer of FBs at the middle- and late-advanced stages was still clearly visible, but its cells were much thinner. The cells of the inner lining? also became thin. The latter still formed a continuous layer in the FBs at the middle-advanced stage of degradation (Fig. 10B), while only a few of these cells survived to the late-advanced stage (Fig. 10C). Cells of the inner layer greatly decreased in size and often had an electron-dense cytoplasm with a still-recognizable RER. Some of them still possessed reduced cytoplasmic processes protruding into the internal cavity (Figs 8B, C, 9D, 10C).

**Figure 9.**
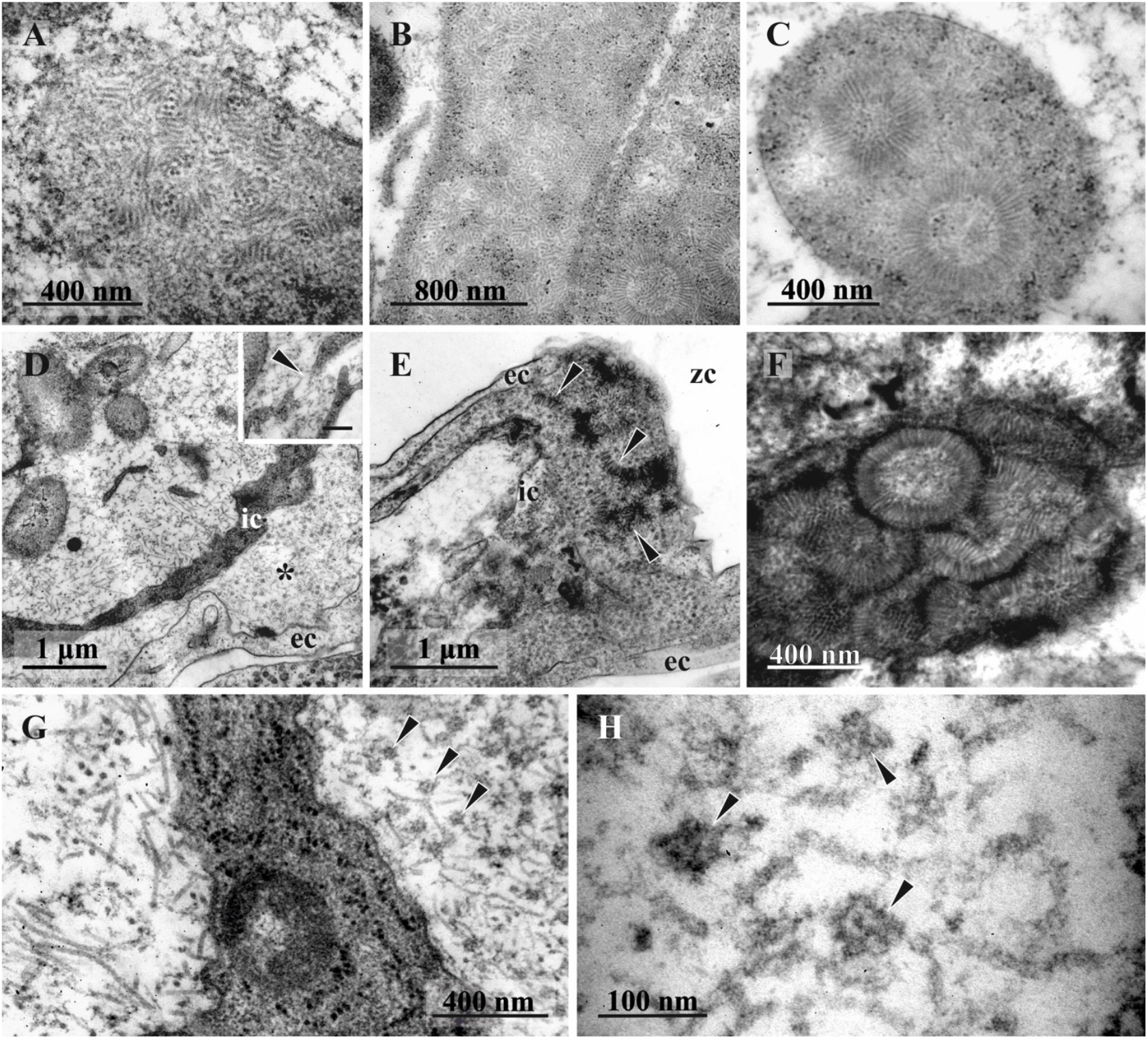
Stages of formation of putative virus-like particles in the funicular bodies at late-advanced stages of degradation (collected on 23 August 2021 and 31 September 2019) (TEM). A, B, C, VLP and ‘self-constructed’ spherical complexes inside bacteria (23 August 2021); D, VLP in symbiont-containing cavity near the bacteria and in ILS between two FB cell layers (asterisk) (inset: probable perforation of the inner cell layer acting as a passage for VLPs transfer to ILS shown by arrowhead, scale bar 200 nm) (23 August 2021); E, F, Self-assembling/constructing of spherical complexes (arrows) inside ILS (31 September 2019). G, ‘Globules’ (arrowheads) and ‘fibrils’ inside ILS (to the right), and filaments in FB inner cavity (to the left) divided by inner layer cell; H, ‘globules’ (arrowheads) and ‘fibrils’ inside ILS under higher magnification. Abbreviations: ec, cells of external layer; ic, cells of inner layer; zc, zooidal cavity.

**Figure 10.**
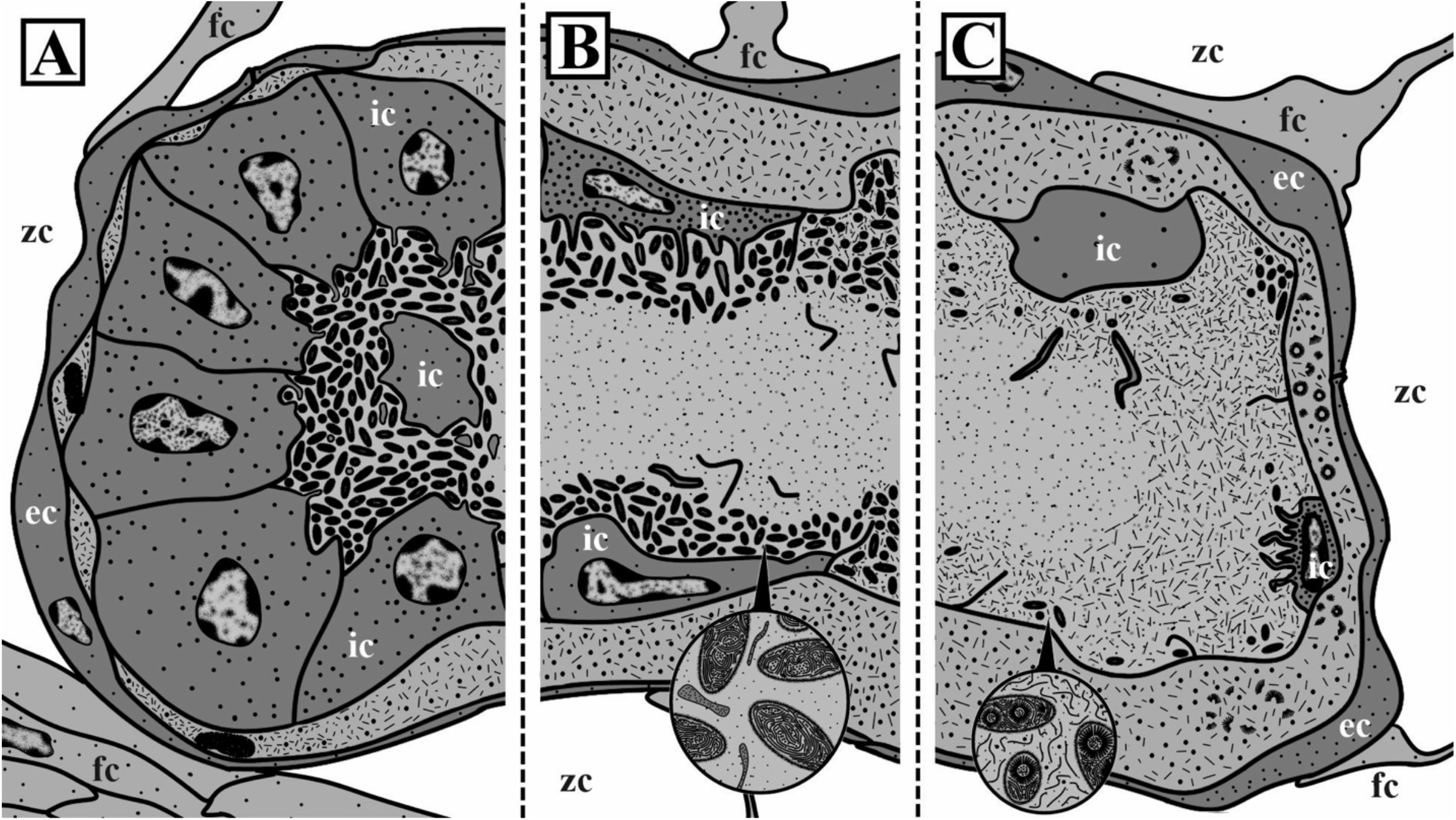
Schemes of funicular bodies of *Dendrobeania fruticosa* at various stages of degradation (from left to right: early-, middle- and late-advanced), showing changes in their structure accompanied by bacterial degradation and putative VLP appearance (magnified sector in B) and self-construction of spherical complexes inside bacteria (magnified sector in C) and inside ILS. Note continuous ‘double’ membrane between internal cavity of funicular body and intercellular space, remaining after collapse of the inner cell layer. Abbreviations: ec, cells of external layer; fc, funicular cell; ic, cells of inner layer; zc, zooidal cavity.

Finally, the collapsed cells of the inner cell layer degraded, leaving only two membranes that ‘stuck’ together. Thus, the internal cavity of late-advanced FBs was enveloped by these membranes and the few remaining inner cells (Figs 8B, 10B, C). In both middle- and late-advanced stages, some bacteria were still visible on the periphery of the FB internal cavity (Figs 8, 9D, 10B, C). In addition, this cavity was filled with dense flocculent material and fragments of cell membranes (Figs 8, 10B, C). The remaining bacteria showed further signs of degradation.

Many were swollen and/or irregularly shaped. The nucleoid underwent further fragmentation, and electron-lucent areas appeared inside some bacteria. ILS was wide and prominent. It still possessed some electron-dense bodies (Fig. 8B), although those found in August were smaller than in those in June. Most of the ILS were filled with the presumed virus-like particles (see below).

Only occasional bacteria were recorded in the large swollen FBs at the terminal stage of degradation in old zooids without polypides in the basal part of the colony collected on 14 June, 2021. The inner cell layer was degraded to a double membrane. ILS was greatly enlarged and contained spherical complexes made of putative VLPs (see below). No bacteria and spherical complexes were detected in the terminal stage FBs in the colony collected on 21 March, 2022.

### Putative VLPs in bacterial cells and FBs

In the cytoplasm of some bacteria in FBs at the middle- and late-advanced stages of degradation, our TEM examination revealed numerous filaments (presumed VLPs). These filaments were either chaotically distributed or assembled in rows, stacks or clumps, sometimes forming spheres 400 to 550 nm in diameter (Fig. 9A-C). The wall of such spherical complexes consisted of 70-110 nm long and 10-15 nm wide filaments. For the first time, spheres were detected inside bacteria in FB at the late-advanced stage of degradation in the colony collected on 23 August 2021. Numerous filaments (some up to 450 nm long and 10-15 nm wide) were also detected, lying freely inside the internal cavity of some FBs (Fig. 9D). Sometimes they were also visible inside ILS, possibly entering it through the gaps in the degrading inner cell layer.

ILS also contained putative VLPs, but their structure was different from those found in the FB cavity and bacteria. Tiny fibrils 100-110 nm long were first detected in this space at the initial stage of degradation of FB in the colony collected on 23 June 2018 (Fig. 7F), but they were especially numerous in FBs at the middle- and late stages of degradation in August and September. Compared with filaments inside the internal cavity of FBs, fibrils were shorter, somewhat thinner, and looked crumpled (compare left and right sides of Fig. 9G). We also recorded numerous, often tightly packed, dark angular globules 40-45 nm in diameter inside ILS (Figs 8B, inset, 9D, E, G), visible as penta- or hexahedrons under high magnification (Fig. 9H). The mixture of these globules and aforementioned fibrils was often chaotic (Figs 9D, 10B, C).

Noteworthy, scattered globules and fibrils sometimes also occurred inside the FB internal cavity. Finally, some late-advanced and terminal stage FBs (collected on 31 September 2019 and 14 June 2021, respectively) contained complete or partial spherical complexes inside their ILS (Fig. 9E, F). Their structure and size were similar to those recorded in bacteria, but each sphere had a thin peripheral (and more electron-dense) zone over the distal ends of filaments (90-100 nm long) forming the rest of the sphere ‘wall’. The central part of the sphere was filled with flocculent material. Spheres in the different stages of self-assembly were always seen in groups. Note that the early stage of the grouping of ‘fibrils’ in ILS was already detected in one FB at the initial stage of its degradation (Fig. 7F).

## Discussion

### Funicular bodies: structure, function, and development

The ultrastructural and functional complexity of funicular bodies in bryozoans ^25,46,47^ (our data), as well as the reduction of the genome in their symbiotic bacteria (e.g., *Bugula neritina* ^48^), point at a long-term co-evolution between these organisms, also suggesting that infected bryozoan colonies spend a significant part of their energy budget supporting numerous bacteria inside FBs. All known prokaryote symbionts are apparently non-pathogenic for bryozoans ^49,62^. Instead, the overall evidence indicates a mutualistic relationship. Symbiotic bacteria are known to produce toxic substances (bryostatins in *Bugula* ^63^; bryoanthratiophene in *Watersipora* ^64^) that protect bryozoan larvae from predators ^42,44,45^. Another potential role of bacterial secondary metabolites is the chemical defense of early developmental stages during larval settlement and metamorphosis ^40,65^. Such chemical defence may also prevent epibiotic overgrowth of the bryozoan colonies by other bacteria and algae (reviewed in ^66,67^). Similar functions can be assumed for symbiotic prokaryotes of *Dendrobeania fruticosa*, although experimental evidence is still required.

FBs with bacterial symbionts in colonies of *D. fruticosa* show signs of high specialization. The walls of the funicular body completely isolate its internal cavity from the surrounding zooidal cavity: the cells of the outer layer overlap each other, whereas cells of the inner layer have tight Z-shaped contacts. Such isolation probably creates a specific environment inside the symbiont-containing space that helps maintain a growing population of bacterial symbionts. A massive protein-synthesizing apparatus was observed in the cells of the inner layer. In addition, the ‘pocket’-like structure of the internal cavity of an FB and the abundance of cytoplasmic processes, some of which protruded deep into this cavity, contribute to increasing both the general inner surface area of the FB and the contact area between its inner cells and bacteria. From their side, bacteria provide this contact by numerous pili. Numerous pits and microvesicles associated with the apical membrane of the inner cells imply an active exchange of substances between the host and its symbionts (e.g., nutrition provided to growing and multiplying bacteria and absorption of the potential wastes they produce).

The ultrastructure and functional morphology of FBs in *Aquiloniella scabra*, the only species in which the ultrastructure of these organs was described, are very similar to those in *D. fruticosa*. Its FBs also consist of two cell types, but the inner cell layer is formed by one or a few cells, whereas external cells form a multilayered envelope ^25^. In both species, the main function of FBs was considered to be the chambers/organs for symbiont incubation and nourishment.

FBs with symbiotic bacteria have been found in several bryozoan species from four different families ^20,25,47,49,51^ (our data). Remarkably, all these families (Bugulidae, Beaniidae, Candidae, Epistomiidae) belong to the clade Buguloidea. Encapsulated aggregations of bacteria within the zooidal cavity were also described in two species of *Watersipora* from the phylogenetically distant family Watersiporidae (Smittinoidea) ^62,68^. In that case, prokaryotes (“mollicutes”) were also enveloped by flattened bryozoan cells, although the current scarcity of data makes it difficult to compare these cell aggregations with funicular bodies.

Among Buguloidea, FBs show the same basic structure, although species described by Lutaud ^49^, Dyrynda and King ^51^, and Mathew with co-authors ^46^ based on light microscopy require reexamination by TEM. Moreover, FBs are in all instances associated with the funicular system of the colony. This similarity could indicate the single origin of the bacterial symbiosis and FBs within Buguloidea. Still, it remains unknown (although rather probable) whether that system transports nutrients from feeding polypides to FBs, because no communication between the lacunae of the funicular cords and the FB internal cavity was detected. Another, more likely option, however, is the independent acquisition of bacteria (see also below) and a similar ‘reaction’ of host tissues to invaders, resulting in the formation of bacterial organs (FB) with a similar bauplan. The presence of incapsulated bacteria in non-related *Watersipora* supports the second interpretation. Answering this question will require both ultrastructural studies and molecular identification of bacteria.

As for the initial source of somatic cells and development of FB, two variants have been suggested. Describing the development of FBs in *Bugulina turbinata*, Lutaud ^49^ stated that the epithelial cells of the cystid wall were transformed into the inner cell layer of the FB, whereas peritoneal cells formed its external lining. By contrast, while studying *Aquiloniella scabra*, Karagodina with co-authors ^25^ suggested that bacteria are engulfed by one of the funicular cells, which becomes a ‘bacteriocyte’ that is later enveloped by neighboring funicular cells. In the latter case, FBs are considered to be modified expanded parts of the funicular system. This is consistent with experiments by Sharp and co-authors ^40^, who detected groups of labeled bacteria in the funicular cords of *Bugula neritina*, and also multiple bacteria developing inside enlarged funicular cords (in fact, very large FBs) in the related *Paralicornia sinuosa* ^20^.

Our TEM study of FBs in *D. fruticosa* showed that they are not swollen parts of the funicular cords, as was stated by Vishnyakov with co-authors for *B. neritina* ^20^. It is more likely that the funicular cords and processes of their cells contact the external cell layer of FBs.

According to the third scenario that we present here, the inner cell layer of FBs in *D. fruticosa*, as well as in other studied bugulids, most likely originates from the coelomocytes which accumulate bacteria via phagocytosis. Such solitary cells were described inside the zooidal cavity in *B. neritina* ^20^. Instead of being digested, engulfed bacteria could trigger the coelomocyte divisions resulting in the formation of the inner cell layer. In contrast, in *A. scabra*, the coelomocyte can remain single or undergo only a few divisions. We propose that the external cell layer of FB in *D. fruticosa* originates from the funicular cells because these cell types are ultrastructurally similar. Finally, it is also possible that the exact process and sources of FB formation differ in different species.

Multiple and lobed FBs found in two zooids of *D. fruticosa* could indicate a potential mode of their multiplication. The case of *P. sinuosa* requires additional study, but currently we believe that its bacteria-bearing ‘funicular cords’ are very large, elongated FBs, as well (see ^20^).

### Symbiont circulation in the bryozoan life cycle

The taxonomic diversity of bryozoan hosts and their symbiotic bacteria – supported by a variety of sites in the bryozoan zooids and larvae where symbionts have been found – unambiguously point to multiple independent origins of symbiotic associations between bacteria and cheilostome Bryozoa ^20,25^.

Rod-shaped bacteria are the most common symbionts in the superfamily Buguloidea. Although superficially similar, these bacteria strongly differ in their maximum size, suggesting the presence of different procaryote species. Thus, bacteria detected in the larvae of *Bugulina simplex* and in FBs of *Aquiloniella scabra* can reach 10 μm in length ^25,69^, while healthy symbionts (see below) inside coelomocytes and presumably peritoneal cells of *Bugula neritina* were only 2.5 μm long ^20^. The maximum length of bacteria in FBs of *D. fruticosa* never exceeded 5 μm. Coccoid bacteria in the tentacles of *B. neritina* were 0.5-0.7 μm in diameter ^20^. Else, oval or irregularly-shaped mycoplasma-like α-Proteobacteria were detected in the genus *Watersipora* ^62,68,720^. By contrast, the symbionts identified in *B. neritina* and *B. simplex* belong to γ-Proteobacteria ^69,71,72^.

Apart from FBs, prokaryote symbionts were described extracellularly in colonies of different cheilostome species: in vestibular glands of autozooids, inside polymorphic zooids (avicularia) ^38,39,50^, in tentacles and funicular cords ^20^. They have also been found intracellularly: inside coelomocytes, epithelial and peritoneal cells of the body wall, and pharyngeal cells ^20,50^. In addition, Woollacott and Zimmer ^73^ described bacteria in the ‘channels’ of the funicular cords associated with brood chambers. However, the TEM image they published shows bacteria inside large vacuoles of the funicular cells – seemingly not in the lacunae between these cells, recalling the aforementioned idea of a ‘bacteriocyte’. Finally, bacteria have also been found in the pallial sinus of bryozoan larvae (^62,68,74^; reviewed in ^72^). All these diverse data have led to two opposite views on the acquisition and circulation of symbionts in the bryozoan life cycle.

Discovery of bacteria in both colonies and larvae of *B. neritina* was regarded as possible evidence of their transmission from larvae to adults ^74^. This assumption was experimentally proven using both labeled bacteria and their metabolites (bryostatins) in the larvae and preancestrulae developing from them ^40^. Moreover, the presence of bacteria within brood chambers (ovicells) in this species was considered as proof for the next step – the transition of symbionts from the colony to the incubated larvae, i.e., the vertical transfer of symbionts. Symbiotic bacteria populating larvae (and making them unpalatable for predators) are incorporated into the preancestrular tissues during larval metamorphosis, and then found inside zooidal buds in early colonies (a symbiont association with the host cells was not specified) and funicular cords of rhizoids in adult colonies ^40^.

The next step of the bacterial development could be the formation of FBs as a locus of symbiont reproduction. Mature FBs, full of bacteria, were considered to be the starting point for the transfer of prokaryotes (by an unknown mechanism) from the zooid to the brood cavity (via funicular cords associated with both FB and ovicells), and then to the incubated larvae ^47^. Light microscopic data demonstrated: (1) the association of FBs with tube-like funicular cords ^47^, and (2) the presence of groups of ‘bacterial bodies’ (small aggregations of bacteria) inside the ooecial vesicle (membranous-epithelial ‘plug’ that closes the entrance to the ovicell; mentioned in *B. neritina* ^47^, and shown in images of the related *Bugulina flabellata* ^75-77^). In addition, ultrastructural data proved the presence of bacteria inside funicular cords, more precisely – inside their funicular cells (see above), “extending to the ooecial vesicle” (^73^, p. 362). Elsewhere, Sharp and co-authors (^40^, p. 697) used fluorescence microscopy to demonstrate the presence of bacteria “within the ovicells”, and suggested that they are transported there across the colony via funicular cords that also house bacteria. Combined, all these data imply that bacteria move from FBs to the ooecial vesicle, accumulate there, and then somehow enter the brood cavity, which contains a larva, either through or in-between the epithelial cells and the cuticle of the ooecial vesicle. Findings of bacteria inside larvae and adult colonies of two species from the non-related genus *Watersipora* ^62,68^ further strengthened the hypothesis of vertical transfer, which has subsequently been widely accepted by many authors ^20,30,47,78^. Despite extensive TEM studies, no bacteria have been found inside the funicular cords in *B. neritina* (Vishnyakov & Ostrovsky, unpublished data), which contradicts the data of Sharp and coauthors ^40^ obtained by fluorescent microscopy Accordingly, it was suggested that coelomocytes carry symbionts to the ooecial vesicle instead ^20^.

Nonetheless, the hypothesis of vertical transfer faces serious objections based on life history, molecular and morphological data. In *Dendrobeania fruticosa* in the White Sea, for example, larval production occurs predominantly in autumn (mainly in the distal parts of branches, Fig. 1A), and no bacteria are present in colonies during this period. In addition, molecular population studies revealed that *B. neritina* is a complex of sibling species, both symbiotic and aposymbiotic, some of which live in sympatry, with the horizontal transfer between colonies being the most parsimonious explanation for the distribution of bacteria between siblings ^79^. A study of the genome of symbiotic bacteria showed that they may be able to live outside the host ^48^, which is consistent with the hypothesis of horizontal transfer (which is not the same as environmental transmission, see below).

TEM data showed no communication between the FB cavity and lacunae of the funicular cords in the studied species, in particular in *B. neritina* (Vishnyakov & Ostrovsky, unpublished data), *Aquiloniella scabra* ^25^ and *D. fruticosa* (this study). Coelomocytes (and presumed peritoneal cells) with bacteria embedded in their cytoplasm were indeed recorded inside the ooecial vesicle in *B. neritina* ^20^. Nevertheless, extensive TEM studies of *B. neritina* ovicells at various stages of placental development (Vishnyakov & Ostrovsky, unpublished data) have not revealed bacteria between placental cells adjoining a developing embryo. The fact that these cells are provided with both tight and adherens junctions (e.g., *B. neritina* and *Bicellariella ciliata* ^73,80^), and additionally are covered by a cuticle (albeit thin) raises the question of whether coelomocytes with bacteria and/or bacteria alone can move through the very thick hypertrophied placental epithelium. Interestingly, Miller with co-authors ^48^ detected a gene encoding chitinase in the genome of the symbiont of *B. neritina* that could potentially be used for cuticle piercing.

Another opportunity for the vertical transfer of symbionts is their transport via the supraneural coelomopore during oviposition (see ^20,81^). In this case, free bacteria in the cavity of the maternal zooid could stick to the ovulated oocyte before its transfer into the brood cavity via the coelomopore. However, free bacteria were never recorded in the zooidal cavity. For the *Watersipora* species, an assumed variant of symbiont transmission is through a strand of mucus extending from the maternal zooid to the released larva and tethering it for a few minutes ^68^.

Environmental transmission, when bacteria are acquired from the surrounding seawater, is an alternative option for symbiont acquisition. It may potentially occur either via infection of brooded larvae inside the ovicell by bacteria entering the brood cavity from the external environment, or via infection of larvae during the free-swimming period by bacteria from the water column. Published images by Sharp and co-authors ^40^ showed the presence of both symbiotic bacteria and bryostatins both inside the ooecial vesicle and in the peripheral part of the brood cavity, close to the entrance of the ovicell. Although these authors stated that such close “locations of both the bacterial symbionts and the bryostatins demonstrate that the *B. neritina*-’*E. sertula*’ association has a delivery system for both the symbionts and the bryostatins to embryos within the ovicell” (^40^, p. 699), we argue, based on the above-mentioned data, that this statement remains a probable yet unproven speculation. Our numerous unpublished TEM images indicate the presence of a large number of bacteria filling the brood space between the embryos and the ovicell wall in the brood chambers with and without developing larvae. These bacteria, attracted by some chemical signal(s), can enter the ovicells from outside and infect larvae. Until the transfer of bacteria through the wall of the ooecial vesicle or during oviposition is documented, environmental transmission remains the more probable method. Notably, recent studies on sponge microbiotas showed that the environmental transmission is widespread in this group of suspension feeders ^82^.

The hypothesis of external acquisition of bacteria by the bryozoan hosts leaves different infection pathways open. Some of these could potentially develop into vertical transfer. Beyond the infection of larvae, prokaryotes could enter feeding autozooids via the mouth (and further through the intestinal epithelium into the zooidal cavity), through the coelomopore — a presumed entrance for alien sperm ^76,81,83^, or by direct infection of the tentacles (probably by penetration through the outer epithelium).

We should stress that FBs were absent in zooidal buds and the youngest zooids with functional polypides in the growing branch tip of one colony of *D. fruticosa* collected in June. This suggests that bacteria are not transmitted from the older colony parts (and, thus, are not inherited from the founding larvae), but obtained from the external medium since older (and more proximal) zooids had FBs. This idea is supported by the lack of signs of transfer of bacteria between zooids via communication pores and their pore-cell complexes in *Dendrobeania fruticosa* and *Aquiloniella scabra* in our TEMs. In contrast, fluorescence microscopy showed symbionts in the non-feeding zooidal bud of the newly-formed small colony of *B. neritina* ^40^. Bacteria were also present in the preancestrula formed during larval metamorphosis. It remains unknown whether they can move from the preancestrula to the bud along with coelomocytes, with growing funicular cords or both before the formation of transverse walls that isolate newly budded zooids. So, interzooidal transfer to budding sites is possible, and the youngest zooids with functional polypides in the growing tips of *D. fruticosa* could already receive bacteria too, but FBs were not yet developed. Thus the question of interzooidal/intracolonial transport of bacteria remains open.

Two infection pathways – via larvae and by direct penetration through the tissues of the functional polypide, potentially exist in the same species. This is the case in *B. neritina*, which has morphologically different symbionts in FBs and in the tentacles ^20^. Bacteria in the epithelial wall of the tentacle sheath and ooecial vesicle (see above) could potentially get there via both pathways, or enter the zooid via the coelomopore (third way), subsequently becoming entrapped by coelomocytes or cells of the cystid wall.

Finally, the presence of bryozoan sibling species, some of which have symbionts while others do not, and the presence of symbiotic and aposymbiotic colonies within the same species ^79^, suggests that bacteria can be lost and acquired anew at both short- and long-term time scales, as occurs in hermatypic corals and their symbiotic zooxanthellae ^84,85^. In this light, it would be important to know whether FBs can develop anew in *D. fruticosa* after overwintering in the same zooids, or whether they appear only in newly budding zooids. What is the source of bacteria in overwintered colonies? Is it an external infection, or some ‘survivor’-cells (descendants of the bacterial pool from the larva that overwintered inside epithelial cells and/or coelomocytes), or both? This will require further study.

### Symbiont population dynamics in Dendrobeania fruticosa and its potential drivers

Whatever the route used by bacteria to enter zooids, they are apparently immediately ‘trapped’ by somatic cells. Free bacteria have never been observed inside the zooidal cavity, another argument against their passage through the wall of the ooecial vesicle to the ovicell.

We have shown that regardless of the as yet unknown mode of FB development, these temporary organs and the symbionts inside them undergo seasonal changes. Early and mature FBs with non-modified morphology and ‘healthy’ bacteria were found in young zooids only in the colonies collected in June. In one of these colonies, older zooids contained FBs at the initial stage of degradation. In the same month, one colony possessed FBs either at the early-advanced degradation stage (in young zooids) still containing numerous bacteria, or at the late-advanced and even terminal stages in old, presumably overwintered zooids. At the initial stage of modification, the slightly developed ‘interlayer’ space between the inner and outer FB cell layers contained fibrils (presumed virus-like particles). Interestingly, the cells of the inner lining apparently engulfed some of the bacterial cells by phagocytosis, supporting our interpretation of the origin of these cells from coelomocytes in this species.

One to two months later (August-September), all examined FBs were either at the middle-, or the late-advanced stages of degradation. The number of bacteria in the FB internal cavity distinctly decreased, the inner layer of cells became thinner, and in some regions remained only as a double membrane. The protein-synthesis apparatus was seen only occasionally, and engulfed bacteria were no longer visible inside the inner cells. A wide ILS, formed between the cells of the outer and inner layers, contained abundant putative virus-like particles (Figs 2, 10). All these FBs were recorded in feeding and non-feeding zooids in the non-growing distal parts of colony branches.

Polypide recycling and a seasonal drop of planktonic food alone cannot explain these changes. Firstly, modified FBs were found in zooids with both degenerated and functioning polypides. Secondly, the initial stages of FBs degradation were detected in June when phytoplankton is abundant in the White Sea (e.g., ^86^). We therefore propose the following scenario for the sequence of changes in FBs and their possible causes. In June, newly-formed zooids build funicular bodies containing bacteria that were acquired either from outside or via internal transfer from older colony parts. During that month, FBs begin to degrade. This process continues throughout the rest of the summer. The mid-advanced stage of FB degradation, with few modified bacteria surviving and distributed on the periphery of the FB cavity, were recorded in August. This stage is reminiscent of the final stage in Lutaud’s ^49^ descriptions of the gradual destruction of FBs accompanied by the disappearance of bacterial symbionts in *Bugulina turbinata*. In late September, FBs change and bacteria disappear, probably through viral lysis (the induction of prophages) (Fig. 2). Young and non-modified FBs were never encountered in August and September, indicating that development of FBs occurs only in young zooids at the growing tips of colony branches in June.

Vishnyakov and co-authors ^20^ recently described the degradation of symbiotic prokaryotes in *Bugula neritina* and *Paralicornia sinuosa* accompanied by a change in bacterial morphology similar to bacteriophage-mediated lysis. Degradation process was accompanied by the appearance of polyhedral VLPs in *B. neritina* and by the formation of structures similar to the so-called metamorphosis-associated contractile complexes (MACs) in *P. sinuosa*. These complexes are phage-related structures whose activity eventually results in the cell lysis, see ^87^. Although the fate of FBs was not analyzed in their paper, these two VLP variants were observed both inside the bacteria and in the FB internal space by Vishnyakov and co-authors ^20^.

In *D. fruticosa*, presumed VLPs in the form of spherical complexes (as clusters of straight filaments) are present inside bacteria and, together with their fragments, inside the ILS. The filaments were also observed in the free state, frequently curved (apparently flexible) in the internal cavity of FBs. ILS was mostly filled with ‘fibrils’ (potentially representing modified/corrupted filaments) and ‘globules’, although filaments were incidentally recorded inside ILS too. We suggest that, in *D. fruticosa*, filaments may represent bacteriophage virions. This interpretation is supported by their appearance being associated with the degradation of bacterial cells, as in the case of VLP in *B. neritina* and *P. sinuosa*. Nonetheless, the morphology of the spherical complexes built from the filaments in *D. fruticosa* is unique: they do not resemble any known group of bacterial viruses. We found these putative VLPs inside ILS between the outer and inner FB cell layers. It remains unclear whether they travel there from the FB internal cavity or self-assemble inside ILS from individual filaments that were incidentally met there too. We add that spherical complexes, complete or partial, were recorded inside ILS of the funicular bodies in non-overwintered (collected on 31 September, 2019) and presumably overwintered (14 June, 2021) colonies.

A filamentous morphology is known from only one bacteriophage from the order Tubulavirales, which includes two families Inoviridae and Plectoviridae ^88^. Although inoviruses or plectoviruses have never been reported to assemble in any regular macrocomplexes, the ability of the filamentous Pf phages (inoviruses) of *Pseudomonas aeruginosa* to form nearly regular liquid crystalline assemblages was recently demonstrated ^89^ (see also review ^90^). Interestingly the formation of such crystals required interactions with bacterial or eukaryotic polymeric molecules such as polysaccharides, DNA and probably mucin ^89,90^.

The development of filamentous phages in bacterial cells usually does not kill the cells because these viruses assemble, along with extrusion from the infected bacterium, without disrupting its cell wall (reviewed in ^91^). However, cell death mediated by filamentous prophage induction has been reported in *P. aeruginosa* due to the emergence of so-called superinfective phage variants ^92-94^. Accordingly, the degradation of bacteria in *D. fruticosa* associated with bacterial viruses is possible.

Since the assembly of known filamentous phages is associated with their extrusion from the cell, virions should not accumulate inside bacteria. Although filamentous assembly intermediates may be present (see ^91^), they are not expected to accumulate in such large quantities and/or form superstructures in the bacterial cytoplasm like the ones we found in *D. fruticosa* (Fig. 9). Our observations revealed nothing resembling the extrusion of a filamentous phage from the surface of a bacterial cell. Therefore, if the described filaments are indeed VLPs, they may represent a new type of bacterial virus.

Even though many details remain unknown, we assume that the filaments, ‘globules’, ‘fibrils’, and spherical structures in *D. fruticosa* are of viral origin. Their development following the total disappearance of bacteria in FBs indicates their bacteriophage nature. If so, our observations support the idea that viruses control the number of symbionts in their bryozoan host ^20^, as has been reported in some insects ^85,95^.

## Acknowledgments

This study was conducted using facilities and equipment of the Educational and Research Station “Belomorskaia”, the Department of Invertebrate Zoology and Resource Centers “Development of Molecular and Cell Technologies” and “Microscopy and Microanalysis”, all Saint Petersburg State University. We thank Mr. S.V. Bagrov and Dr N.N. Shunatova, Department of Invertebrate Zoology, Saint Petersburg State University, for their help in collecting the material. We thank Dr. M. Stachowitsch, University of Vienna, for linguistically revising the early draft of the manuscript. Four reviewers critically assessed the MS and gave valuable comments for its improvement.

## Authors’ Contributions

A.O. and A.V. designed the study. O.K. collected material. E.B. and A.V. performed microscopical study. E.B. and A.O. analyzed the data and wrote the manuscript. A.V., A.L., O.K. and A.G. contributed to data interpretation and writing the manuscript. All authors have read and approved the manuscript.

## Ethics approval

No ethical issues were raised during our research.

## Sampling and field studies

No permits for sampling and field studies were required.

## Data availability

All the data produced during this study are available from Andrew Ostrovsky.

## References

1. Dogiel, V. A. Parasitology (Leningrad University Press, Saint Petersburg, 1962). [In Russian]

2. Boucher, D. H., James, S. & Keeler, K. H. The ecology of mutualism. Annu. Rev. Ecol. Syst. 13, 315–347 (1982).

3. Gilbert, S. F., Sapp, J. & Tauber, A. I. A symbiotic view of life: We have never been individuals. Q. Rev. Biol. 87, 325–341 (2012).

4. Van Dover, C. The ecology of deep-sea hydrothermal vents. (Princeton University Press, 2000).

5. Taylor, J. D. & Glover, E. A. Chemosymbiotic bivalves. In The vent and seep biota — from Microbes to Ecosystems (ed Kiel, S.) 107–135 (Springer, Heidelberg, 2010).

6. Sorokan, A. V., Burkhanova, G. F., Benkovskaya, G. V. & Maksimov, I. V. Colorado potato beetle microsymbiont Enterobacter BC-8 inhibits defense mechanisms of potato plants using crosstalk between jasmonate-and salicylate-mediated signaling pathways. Arthropod-Plant Int. 14, 161–168 (2019).

7. Bogdanov, E. A., Vishnyakov, A. E. & Ostrovsky, A. N. From Procaryota to Eumetazoa: Symbiotic associations in fossil and recent bryozoans. Paleontol. J. (in press).

8. Johnson, M. D. The acquisition of phototrophy: Adaptive strategies of hosting endosymbionts and organelles. Photosynth. Res. 107, 117–132 (2011).

9. Balzano, S. et al. Transcriptome analyses to investigate symbiotic relationships between marine protists. Front. Microbiol. 6, 98 (2015).

10. Glynn, P. W. Defense by symbiotic Crustacea of host corals elicited by chemical cues from predator. Oecologia 47, 287–290 (1980).

11. Overstreet, R. M. Metazoan symbionts of crustaceans. In The biology of crustacea: Pathobiology, 6 (ed Bliss, D. E.) 155–250 (Academic Press, New York, London, 1983).

12. Ross, A. & Newman, W. A. A new sessile barnacle symbiotic with bryozoans from Madagascar and Mauritius (Cirripedia: Balanomorpha): A unique case of co-evolution? Invert. Biol. 115, 150–161 (1996).

13. Melo Clavijo, J., Donath, A., Serôdio, J. & Christa, G. Polymorphic adaptations in metazoans to establish and maintain photosymbioses. Biol. Rev. 93, 2006–2020 (2018).

14. Bass, D. et al. Parasites, pathogens, and other symbionts of copepods. Trends Parasitol. 37, 875–889 (2021).

15. Gast, R. J., Sanders, R. W. & Caron, D. A. Ecological strategies of protists and their symbiotic relationships with prokaryotic microbes. Trends Microbiol. 17, 563–569 (2009).

16. Decelle, J., Colin, S. & Foster, R. A. Photosymbiosis in marine planktonic protists. In Marine protists (eds Ohtsuka, S., Suzaki, T., Horiguchi, T., Suzuki, N. & Not, F.) 465–500 (Springer, Tokyo, 2015).

17. Felbeck, H. & Distel, D. L. Prokaryotic symbionts of marine invertebrates. In: The prokaryotes 3891–3906 (Springer, New York, 1992).

18. Vishnyakov, A. E. & Ereskovsky, A. V. Bacterial symbionts as an additional cytological marker for identification of sponges without a skeleton. Mar. Biol. 156, 1625–1632 (2009).

19. Mohamed, N., Saito, K., Tal, Y. & Hill, R. T. Diversity of aerobic and anaerobic ammonia-oxidizing bacteria in marine sponges. ISME J. 4, 38–48 (2010).

20. Vishnyakov, A. E. et al. First evidence of virus-like particles in the bacterial symbionts of Bryozoa. Sci. Rep. 11, 4 (2021).

21. Trautman, D. A. & Hinde, R. Sponge/algal symbioses: A diversity of associations. In Symbiosis. Cellular origin, life in extreme habitats and astrobiology, vol 4 (ed Seckbach, J.) 521–537 (Springer, Dordrecht, 2001).

22. Baker, A. C. Flexibility and specificity in coral-algal symbiosis: Diversity, ecology, and biogeography of Symbiodinium. Annu. Rev. Ecol. Evol. Syst. 34, 661–689 (2003).

23. Taylor, M. W., Radax, R., Steger, D. & Wagner, M. Sponge-associated microorganisms: Evolution, ecology, and biotechnological potential. Microbiol. Mol. Biol. Rev. 71, 295–347 (2007).

24. Muller-Parker, G., D’Elia, C. F., Cook, C. B. Interactions between corals and their symbiotic algae. In Coral reefs in the Anthropocene. (ed Birkeland, C.) 99–116 (Springer, Dordrecht, 2015).

25. Karagodina, N. P., Vishnyakov, A. E., Kotenko, O. N., Maltseva, A. L. & Ostrovsky, A. N. Ultrastructural evidence for nutritional relationships between a marine colonial invertebrate (Bryozoa) and its bacterial symbionts. Symbiosis 75, 155–164 (2018).

26. McKinney, F. K. & Jackson, J. B. C. Bryozoan evolution. (University of Chicago Press, 1991).

27. Ryland, J. S. Bryozoa: An introductory overview, In Moostiere (Bryozoa) (ed Woess, E.) 16, 9–20 (Denisia, Oberösterreichisches Landesmuseum, 2005).

28. Cook, P., Weaver, H., Bock, P. & Gordon, D. Australian Bryozoa. (CSIRO Publishing, 2018).

29. Mukai, H., Terakado, K. & Reed, C. G. Bryozoa. In Microscopic anatomy of invertebrates. (eds Harrison, F. W., Woollacott, R. M.) 45–206 (Wiley-Liss, Inc, New York, 1997).

30. Schwaha, T. Phylum Bryozoa. In Handbook of Zoology (DeGruyter, Berlin, 2021).

31. Schwaha, T. F., Ostrovsky, A. N. & Wanninger, A. Key novelties in the evolution of the aquatic colonial phylum Bryozoa: Evidence from soft body morphology. Biol. Rev. 95, 696–729 (2020).

32. Nekliudova, U. A. et al. Three in one: evolution of viviparity, coenocytic placenta and polyembryony in cyclostome bryozoans. BMC Ecology and Evolution 21, 1–34 (2021).

33. Serova, K. M. et al. Reduction, rearrangement, fusion and hypertrophy: Evolution of the muscular system in polymorphic zooids of cheilostome Bryozoa. Org. Diver. Evol. (2022).

34. Carle, K. J. & Ruppert, E. E. Comparative ultrastructure of the bryozoan funiculus: A blood vessel homologue. J. Zool. Syst. Evol. Res. 21, 181–193 (1983).

35. Lutaud, G. Preliminary experiments on interzooidal metabolic transfer in anascan bryozoans. In Bryozoa: Ordovician to Recent. (eds Nielsen, C., Larwood, G. P.) 183–191 (Olsen & Olsen, Fredensborg, 1985).

36. Best, M. A. & Thorpe, J. P. Autoradiographic study of feeding and the colonial transport of metabolites in the marine bryozoan Membranipora membranacea. Mar. Biol. 84, 295–300 (1985).

37. Shunatova, N., Denisova, S. & Shchenkov, S. Ultrastructure of rhizoids in the marine bryozoan Dendrobeania fruticosa (Gymnolaemata: Cheilostomata). J. Morphol. 282, 847–862 (2021)

38. Lutaud, G. Sur la structure et le rôle des glandes vestibulaires et sur la nature de certains organes de la cavité cystidienne chez les Bryozoaires chilostomes. Cah. Biol. Mar. 5, 201–231 (1964).

39. Lutaud, G. Sur la presence de microorganismes specifiques dans les glandes vestibulaires et dans laviculaire de Palmicellaria skenei (Ellis et Solander) bryozoaire chilostome. Cah. Biol. Mar. 6, 181–190 (1965).

40. Sharp, K. H., Davidson, S. K. & Haygood, M. G. Localization of ‘Candidatus Endobugula sertula’and the bryostatins throughout the life cycle of the bryozoan Bugula neritina. ISME J. 1, 693–702 (2007).

41. Li, H., Mishra, M., Ding, S. & Miyamoto, M. M. Diversity and dynamics of “Candidatus Endobugula” and other symbiotic bacteria in Chinese populations of the bryozoan, Bugula neritina. Microb. Ecol. 77, 243–256 (2019).

42. Lindquist, N. & Hay, M. E. Palatability and chemical defense of marine invertebrate larvae. Ecol. Monogr. 66, 431–450 (1996).

43. Davidson, S., Allen, S., Lim, G. E., Anderson, C. & Haygood, M. Evidence for the biosynthesis of bryostatins by the bacterial symbiont “Candidatus Endobugula sertula” of the bryozoan Bugula neritina. Appl. Environ. Microbiol. 67, 4531–4537 (2001).

44. Lopanik, N., Gustafson, K. R. & Lindquist, N. Structure of bryostatin 20: A symbiont-produced chemical defense for larvae of the host bryozoan, Bugula neritina. J. Nat. Prod. 67, 1412–1414 (2004).

45. Lopanik, N., Lindquist, N. & Targett, N. Potent cytotoxins produced by a microbial symbiont protect host larvae from predation. Oecologia 139, 131–139 (2004).

46. Mathew, M. et al. Influence of symbiont-produced bioactive natural products on holobiont fitness in the marine bryozoan, Bugula neritina via protein kinase C (PKC). Mar. Biol. 163, 1–17 (2016).

47. Mathew, M., Schwaha, T., Ostrovsky, A. N. & Lopanik, N. B. Symbiont-dependent sexual reproduction in marine colonial invertebrate: Morphological and molecular evidence. Mar. Biol. 165, 14 (2018).

48. Miller, I. J., Vanee, N., Fong, S. S., Lim-Fong, G. E. & Kwan, J. C. Lack of overt genome reduction in the bryostatin-producing bryozoan symbiont “Candidatus Endobugula sertula”. Appl. Environ. Microbiol. 82, 6573–6583 (2016).

49. Lutaud, G. La nature des corps funiculaires des cellularines, bryozoaires chilostomes. Arch. Zool. Exp. Gen. 110, 5–30 (1969).

50. Lutaud, G. L’infestation du myoépithélium de l’oesophage par des microorganismes pigmentés et la structure des organes à bactéries du vestibule chez le Bryozoaire Chilostome Palmicellaria skenei (E. et S.). Can. J. Zool. 64, 1842–1851 (1986).

51. Dyrynda, P. E. J. & King, P. E. Sexual reproduction in Epistomia bursaria (Bryozoa: Cheilostomata), an endozooidal brooder without polypide recycling. J. Zool. 198, 337–352 (1982).

52. Packard, A. S. A list of animals dredged near Caribou Island, southern Labrador, during July and August, 1860. Can. Natur. Geol. 8, 401–429 (1863).

53. Osburn, R. C. Bryozoa of the Pacific Coast of America. Part 1, Cheilostomata–Anasca. Allan. Hancock. Pac. Exped. 14, 1–269 (1950).

54. Kluge, G. A. Bryozoa of the northern seas of the USSR. (Amerind Publishing Co, New Delhi, 1975).

55. Gostilovskaya, M. G. Bryozoa of the White Sea. (Nauka, Leningrad, 1978). [In Russian]

56. Gontar, V. I. Bryozoa of the Laptev Sea and New Siberian shoals. Explorations of the Fauna of the Seas 37, 130–138 (1990). [In Russian with English summary]

57. Ryland, J. S. & Hayward, P. J. Erect Bryozoa. Marine flora and fauna of the northeastern United States. NOAA Technical Report NMFS 99, 1–47 (1991).

58. Grischenko, A. V. Bryozoans (Ctenostomida, Cheilostomida) of the shelf zone of the Commander Islands. In Benthic flora and fauna of the shelf zone of the Commander Islands (ed Rzhavsky, A. V.) 153–192 (Dalnauka, Vladivostok, 1997). [In Russian with English summary]

59. Hayward, P. J. & Ryland, J. S. Cheilostomatous Bryozoa, Part 1, Aeteoidea–Cribrilinoidea. Synopses of the British Fauna (New Series), No. 10. 2nd ed. (The Linnean Society of London and The Estuarine and Brackish-water Sciences Association, London, 1998).

60. Richardson, K. C., Jarett, L. & Finke, E. H. Embedding in epoxy resins for ultrathin sectioning in electron microscopy. Stain Technol. 35, 313–323 (1960).

61. Humphrey, C. D. & Pittman, F. E. A simple methylene blue-azure II-basic fuchsin stain for epoxy-embedded tissue sections. Stain Technol. 49, 9–14 (1974).

62. Boyle, P. J., Maki, J. S. & Mitchell, R. Mollicute identified in novel association with aquatic invertebrate. Curr. Microbiol. 15, 85–89 (1987).

63. Pettit, G. R. et al. Isolation and structure of bryostatin 1. J. Am. Chem. Soc. 104, 6846–6848 (1982).

64. Jeong, S. J. et al. Bryoanthrathiophene, a new antiangiogenic constituent from the bryozoan Watersipora subtorquata (d’Orbigny, 1852). J. Nat. Prod. 65, 1344–1345 (2002).

65. Lopanik, N., Targett, N. & Lindquist, N. Ontogeny of a symbiont-produced chemical defense in Bugula neritina (Bryozoa). Mar. Ecol. Prog. Ser. 327, 183–191 (2006).

66. Sharp, J. H., Winson, M. K. & Porter, J. S. Bryozoan metabolites: An ecological perspective. Nat. Prod. Rep. 24, 659–673 (2007).

67. Winston, J. E. & Migotto, A. E. Behavior. In Phylum Bryozoa. Handbook of zoology (ed Schwaha, T.) 143–187 (DeGruyter, Berlin, 2021).

68. Zimmer, R. L. & Woollacott, R. M. Mycoplasma-like organisms: Occurrence with the larvae and adults of a marine bryozoan. Science 220, 208–210 (1983).

69. Lim, G. E. & Haygood, M. G. “Candidatus Endobugula glebosa,” a specific bacterial symbiont of the marine bryozoan Bugula simplex. Appl. Environ. Microbiol. 70, 4921–4929 (2004).

70. Anderson, C. M. & Haygood, M. G. α-Proteobacterial symbionts of marine bryozoans in the genus Watersipora. Appl. Environ. Microbiol. 73, 303–311 (2007).

71. Haygood, M. G. & Davidson, S. K. Small-subunit rRNA genes and in situ hybridization with oligonucleotides specific for the bacterial symbionts in the larvae of the bryozoan Bugula neritina and proposal of” Candidatus endobugula sertula”. Appl. Environ. Microbiol. 63, 4612– 4616 (1997).

72. Lim-Fong, G. E., Regali, L. A. & Haygood, M. G. Evolutionary relationships of “Candidatus Endobugula” bacterial symbionts and their Bugula bryozoan hosts. Appl. Environ. Microbiol. 74, 3605–3609 (2008).

73. Woollacott, R. M. & Zimmer, R. L. A simplified placenta-like system for the transport of extraembryonic nutrients during embryogenesis of Bugula neritina (Bryozoa). J. Morphol. 147, 355–377 (1975).

74. Woollacott, R. M. Association of bacteria with bryozoan larvae. Mar. Biol. 65, 155–158 (1981).

75. Ostrovsky, A. N., Gordon, D. P. & Lidgard, S. Independent evolution of matrotrophy in the major classes of Bryozoa: Transitions among reproductive patterns and their ecological background. Mar. Ecol. Prog. Ser. 378, 113–124 (2009).

76. Ostrovsky, A. N. Evolution of sexual reproduction in marine invertebrates: Example of gymnolaemate bryozoans. (Springer, Dordrecht, Heidelberg, New York, London, 2013).

77. Ostrovsky, A. N. From incipient to substantial: Evolution of placentotrophy in a phylum of quatic colonial invertebrates. Evolution 67, 1368–1382 (2013).

78. Miller, I. J., Weyna, T. R., Fong, S. S., Lim-Fong, G. E. & Kwan, J. C. Single sample resolution of rare microbial dark matter in a marine invertebrate metagenome. Sci. Rep. 6, 34362 (2016).

79. Linneman, J., Paulus, D., Lim-Fong, G. & Lopanik, N. B. Latitudinal variation of a defensive symbiosis in the Bugula neritina (Bryozoa) sibling species complex. PLOS One 9, e108783 (2014).

80. Moosbrugger, M., Schwaha, T., Walzl, M. G., Obst, M. & Ostrovsky, A. N. The placental analogue and the pattern of sexual reproduction in the cheilostome bryozoan Bicellariella ciliata (Gymnolaemata). Front. Zool. 9, 29 (2012).

81. Ostrovsky, A. N. & Porter, J. S. Pattern of occurrence of supraneural coelomopores and intertentacular organs in Gymnolaemata (Bryozoa) and its evolutionary implications. Zoomorphology 130, 1–15 (2011).

82. Björk, J. R., Díes-Vives, C., Astudillo-Garsia, C., Archie, E. A. & Montoya, J. M. Vertical transmission of sponge microbiota is inconsistent and unfaithful. Nat. Ecol. Evol. 3, 1172–1183 (2019).

83. Reed, C. G. Bryozoa. In Reproduction of marine invertebrates, VI. Echinoderms and Lophophorates (eds Giese, A. C., Pearse, J. S., Pearse, V. B.) 85–245 (Boxwood Press, Pacific Grove, 1991).

84. Toller, W. W., Rowan, R. & Knowlton, N. Repopulation of zooxanthellae in the Caribbean corals Montastraea annularis and M. faveolata following experimental and disease-associated bleaching. Biol. Bull. 201, 360–373 (2001).

85. Bordenstein, S. R., Marshall, M. L., Fry, A. J., Kim, U. & Wernegreen, J. J. The tripartite associations between bacteriophage, Wolbachia, and arthropods. PLoS Pathog. 2, e43 (2006).

85. Coffroth, M. A., Poland, D. M., Petrou, E. L., Brazeau, D. A. & Holmberg, J. C. Environmental symbiont acquisition may not be the solution to warming seas for reef-building corals. PLoS One 5, e13258 (2010).

86. Shevchenko, E. T. et al. Electra vs Callopora: Life histories of two bryozoans with contrasting reproductive strategies in the White Sea. Invertebr. Reprod. Dev. 64, 137–157 (2020).

87. Shikuma, N. J. et al. Marine tubeworm metamorphosis induced by arrays of bacterial phage tail-like structures. Science 343, 529–533 (2014)

88. Koonin, E. V. et al. Global organization and proposed megataxonomy of the virus world. Microbiol. Mol. Biol. Rev. 84, e00061–19 (2020).

89. Secor, P. R. et al. Biofilm assembly becomes crystal clear–filamentous bacteriophage organize the Pseudomonas aeruginosa biofilm matrix into a liquid crystal. Microb. Cell. 3, 49– 52 (2015).

90. Secor, P. R. et al. Pf bacteriophage and their impact on Pseudomonas virulence, mammalian immunity, and chronic infections. Front. Immunol. 11, 244 (2020).

91. Rakonjac, J., Russel, M., Khanum, S., Brooke, S. J. & Rajič, M. Filamentous phage: Structure and biology. Adv. Exp. Med. Biol. 1–20 (2017).

92. Rice, S. A. et al. The biofilm life cycle and virulence of Pseudomonas aeruginosa are dependent on a filamentous prophage. ISME J. 3, 271–282 (2009).

93. Hui, J. G. K., Mai-Prochnow, A., Kjelleberg, S., McDougald, D. & Rice, S. A. Environmental cues and genes involved in establishment of the superinfective Pf4 phage of Pseudomonas aeruginosa. Front. Microbiol. 5, 654 (2014).

94. Tortuel, D. et al. Activation of the cell wall stress response in Pseudomonas aeruginosa infected by a Pf4 phage variant. Microorganisms 8, 1700 (2020).

95. Weldon, S. R.. & Oliver, K. M. Diverse bacteriophage roles in an aphid-bacterial defensive mutualism. In The mechanistic benefits of microbial symbionts (ed Hurst, C. J.) 173–206 (Springer, Cham, 2016).

